# The development and neuronal complexity of bipinnaria larvae of the sea star *Asterias rubens*

**DOI:** 10.1101/2021.01.04.425292

**Authors:** Hugh F. Carter, Jeffrey R. Thompson, Maurice R. Elphick, Paola Oliveri

## Abstract

Free-swimming planktonic larvae are a key stage in the development of many marine phyla, and studies of these organisms have contributed to our understanding of major genetic and evolutionary processes. Although transitory, these larvae often attain a remarkable degree of tissue complexity, with well-defined musculature and nervous systems. Amongst the best studied are larvae belonging to the phylum Echinodermata, but with work largely focused on the pleuteus larvae of sea urchins (class Echinoidea). The greatest diversity of larval strategies amongst echinoderms is found in the class Asteroidea (sea-stars), organisms that are rapidly emerging as experimental systems for genetic and developmental studies. However, the bipinnaria larvae of sea stars have only been studied in detail in a small number of species and the full complexity of the nervous system is, in particular, poorly understood. Here we have analysed embryonic development and bipinnaria larval anatomy in the common North Atlantic sea-star *Asterias rubens*, employing use of a variety of staining methods in combination with confocal microscopy. Importantly, the complexity of the nervous system of bipinnaria larvae was revealed in greater detail than ever before, with identification of at least three centres of neuronal complexity: the anterior apical organ, oral region and ciliary bands. Furthermore, the anatomy of the musculature and sites of cell division in bipinnaria larvae were analysed. Comparisons of developmental progression and molecular anatomy across the Echinodermata provided a basis for hypotheses on the shared evolutionary and developmental processes that have shaped this group of animals. We conclude that bipinnaria larvae appear to be remarkably conserved across ~200 million years of evolutionary time and may represent a strong evolutionary and/or developmental constraint for species utilizing this larval strategy.

## Introduction

Species from many marine phyla develop via a biphasic lifestyle, transitioning from a free-swimming planktonic larva to a more sedentary benthonic adult. Although transitory, the larval phase often develops a remarkable degree of tissue complexity, including a nervous system and musculature. The study of the planktonic larvae of several marine taxa has been important for our understanding of species dispersal (Scheltema 1986) and developmental processes (Raff 2008). Furthermore, since the majority of animal phyla are marine, comparative studies of genetic networks that direct the molecular and cellular processes of larval development are key to our understanding of the evolutionary processes that have shaped the animal kingdom (Davidson et al. 2002; Davidson and Erwin 2006; Peter and Davidson 2015; Dylus et al. 2016).

Of the phyla with planktonic larvae, the Echinodermata is perhaps the best understood, having been used as experimental organisms for studies of early development for over a century (Boveri 1893). Unlike their pentaradial adults, echinoderm larvae are bilaterally symmetrical, although they differ widely in morphology between and within classes (McEdward and Miner 2001; Young et al. 2002). Echinoderm larvae are typically characterised based upon how nutrition is derived prior to metamorphosis: either free-feeding planktotrophs with complex ciliary bands, or lecithotrophs dependent on a maternally derived yolky substance (Zamora et al. 2020). Within these categories there is considerable diversity and some species display highly derived developmental progressions that do not fall neatly within these two sub-divisions (McEdward 1995; Byrne et al. 2007). Although indirect-developing echinoderm larvae vary extensively in morphology within and across classes (Wray 1992; Hart et al. 1997; McEdward and Miner 2001; Byrne and Selvakumaraswamy 2002; Young et al. 2002), phylogenetic comparative studies have concluded that indirect development, through a free-swimming and feeding planktotrophic larva, represents the ancestral strategy for all echinoderms (McEdward and Janies 1997; Peterson et al. 2000; Raff and Byrne 2006). In order to understand conserved and divergent morphologies, tissues, and cell types across echinoderm larvae, a better understanding of their molecular signatures in indirect-developing, planktonic larvae is crucial.

Amongst echinoderms, the Asteroidea or sea stars have the greatest variety of described larval strategies (McEdward and Miner 2001) and are emerging as experimental systems for developmental and genetic studies (Stewart et al. 2015; Byrne et al. 2020; Cary et al. 2020). The bat-star *Patiria miniata* in particular has become an important comparative resource for understanding divergence and conservation of gene regulatory network architecture over long evolutionary timescales (Cary and Hinman 2017). Additionally, a growing number of asteroid species now have well-annotated genome assemblies, making this a particularly pertinent moment for a detailed assessment of asteroid larval development (Hall et al. 2017; Cary et al. 2018; Ruiz-Ramos et al. 2020).

The bipinnaria, a free swimming planktotrophic larvae, is the most phylogenetically widespread larval form amongst asteroids and is considered to represent the ancestral larval form of the class (McEdward 1995; Raff and Byrne 2006). In most cases the bipinnaria is followed by a more complex brachiolaria, from which arises an attachment complex prior to metamorphosis (Haesaerts et al. 2005). Bipinnaria are characterized by two bilaterally symmetrical ciliary bands and an open, functional gut. Although superficially simple, they require a surprising degree of neuronal complexity, both for environmental sensing and for coordination of the ciliary bands, which play a dual role in feeding and locomotion (Burke 1983; Lacalli et al. 1990; Hinman and Burke 2018). Despite its key evolutionary position, the bipinnaria larva has only been the subject of detailed investigation in a small number of species and there remains only a rudimentary understanding of the complexity of its nervous system (Moss et al. 1994; Byrne and Cisternas 2002).

One of the most abundant northern hemisphere asteroids, *Asterias rubens*, populates the Atlantic Ocean from northern Norway to Senegal in the east and northern Canada to North Carolina in the west at depths of 0-650m (Budd 2008). *A. rubens* undergoes the common larval transition of bipinnaria to brachiolaria, but its early larval development from first cleavage to free swimming bipinnaria has not been studied in detail for more than a century (Gemmill 1914). While studies of larval development are lacking, adult *A. rubens* has been an experimental system for functional characterisation of neuropeptide signalling for several decades and a large set of taxon-specific antibodies against multiple neuropeptides has been developed (Elphick et al. 1995; Odekunle et al. 2019; Zhang et al. 2020). This has been facilitated by rapidly increasing availability of transcriptomic and genomic resources for this species (Semmens et al. 2016). The expression patterns of many neuropeptides have been described in adults (Newman et al. 1995; Lin et al. 2017; Tian et al. 2017; Cai et al. 2018) and the brachiolaria stage (Mayorova et al. 2016), allowing for comparison of conserved or differential molecular signatures between the larval and adult body plans. *A. rubens* is also phylogenetically well-suited for comparative studies of the larval nervous system. It belongs to the order Forcipulatida and is thus distantly related to the valvatid *P. miniata*, but closely related to another forcupilatid - *Pisaster ochraeus* (Mah and Foltz 2011). These two species are the asteroids in which the larval nervous system has previously been characterized in detail, and studies on *A. rubens* thus allow for comparisons across both deep and shallow phylogenetic distances.

To precisely understand the development of bipinnaria larvae from a molecular and cellular perspective, and to facilitate evolutionary comparisons across echinoderms and other marine invertebrates, we have surveyed the early and larval development of *A. rubens* by analysing expression of selected proteins and cell proliferation. In particular, we have focused on the larval nervous system through the use of novel antibodies, some of which are here uniquely applied to echinoderm larvae. This work enables comparisons of the development of the nervous system and other structures across the planktonic larvae of the Echinodermata to better provide an understanding of the evolutionary and developmental processes that have shaped the biodiversity of the group.

## Materials and Methods

### Gamete collection and embryo culture

Gamete collection and embryo culture followed Jarvela and Hinman (2014). Following fertilization, eggs were incubated at either 12° or 15°C and embryonic development allowed to proceed (see supplementary methods). Developmental progression was checked hourly until embryos had undergone their fifth cleavage, then at least once daily until the culture reached the bipinnaria stage. At each time point, several individuals (>30) were mounted on glass slides and visualised using light-microscopy. Embryos and larvae were collected at different stages for further processing.

### Immunostaining and cell proliferation

Immunostaining and assays of cell proliferation followed procedures reported by Thompson et al. (2020). For each developmental time point of interest, 4000-7000 embryos or larvae were fixed in 4% PFA, permeabilized in ice-cold methanol and stored in blocking buffer (BB). Samples were incubated in primary antibodies at 37°C for 1.5 hrs, then incubated in secondary antibodies for 1 hr at room temperature (RT) prior to final incubation with DAPI for 5 min at RT. For antibody combinations and further details see supplementary materials. Dividing cell nuclei were labelled using the Click-iT^®^ EdU Alexa Flour^®^ 555HCS kit (Life Technologies) according to the manufacturer’s protocol.

### Microscopy

Differential interference contrast (DIC) and epi-fluorescent images were produced using a Zeiss AxioImager M1 microscope and Zeiss AxioCamHRc camera. Confocal microscopy was performed using a LSM 800 confocal microscope or Leica SPEinv inverted confocal microscope. Image processing and analysis was carried out using ImageJ version 2.0.0.0.

## Results

### Asterias rubens develops canonically to a bipinnaria larvae

To better understand the tempo and mode of *A. rubens* development, we optimized culture conditions to produce thousands of syncronously developing embryos. Development was followed at two different temperatures (12° and 15°C), corresponding to the UK coastal temperature range during the spawning season, in three independent batches to produce a staging system (Fig 1, Supp Fig 1). A full account of larval development can be found in the supplementary material. Briefly, maturation of ooyctes was induced with 1-Methyl-adenine leading to loss of the germinal vesicle (Fig 1A-B). The fertilization membrane elevates rapidly after sperm addition, and first cleavage is complete by 2 hpf at 15°C and 3 hpf at 12°C (Fig 1D). In each batch, subsequent cleavages are holoblastic and equal for roughly 9 to 11 cycles, occurring on average every 2 h at 15°C and 3 h at 12°C. The fertilization membrane constrains growth until ~24 hpf (Fig 1G) with its loss accompanied by a pronounced elongation along the animal/vegetal axis and a thickening of cells in the vegetal half of the embryo (Fig 1H). The vegetal plate flattens and thickens prior to gastrulation (Fig1I), producing an invaginating archenteron that reaches half the length of the embryo by 46 hpf. The lumen of the archenteron gradually decreases in diameter as the walls thicken and the archenteron extends in length (Fig1J-K). By 70 hpf, mesenchyme cells arising from the tip of the archenteron are apparent in the blastocoel (Fig1K) and these bud-off into the blastocoel throughout gastrulation (Fig1K-M). By 92 hpf two differently sized coelomic pouches are apparent and the central part of the archenteron expands to form the proto-stomach (Fig 1L). The immature bipinnaria (122 hpf) has a tripartite gut with an oesophagus, stomach and hind-gut (Fig 1M), and mesenchymal cells that have ingressed into the blastocoel start to show considerable morphological differentiation (Supp Fig 3).

**Figure 1.**
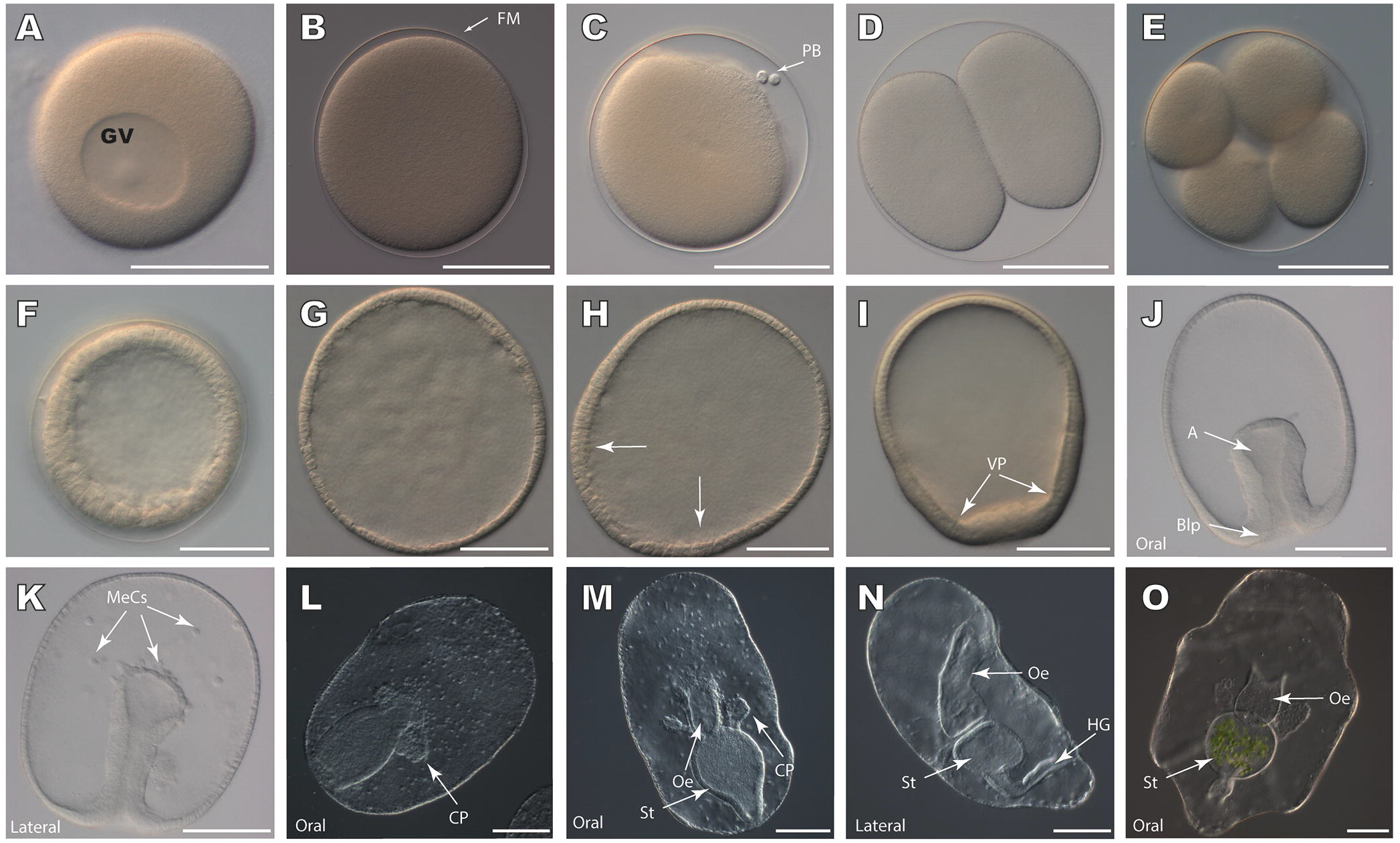
Developmental progression of *A. rubens* from fertilized egg to bipinnaria larva. (A) Unfertilized immature oocyte with germinal vesicle. (B) Fertilized oocyte without germinal vesicle. Note presence of fertilization membrane. (C) Fertilized oocyte with polar bodies. (D) Two-cell stage embryo following the first cleavage. (E) Four-cell stage embryo. Cleavages are roughly equal, giving rise to blastomeres of roughly equal size within the fertilization membrane. (F) Blastula-stage embryo, consisting of approximately 250 cells. Embryos are round at this stage, due to constraints imposed by the fertilization membrane. (G) Blastula-stage embryo following loss of the fertilization membrane. (H) Swimming blastula-stage embryo. (I) Blastula-stage embryo. Elongation has taken place along the antero-posterior axis, and thickening and flattening of the vegetal plate is evident along the vegetal end of the embryo. (J) Early gastrula-stage embryo. Archenteron has invaginated and has thinner walls at the blind end than in the rest of the tube. (K) Mid-gastrula stage embryo. Archenteron has thickened relative to early gastrula-stage embryos seen in (J). Additionally, many mesenchymal cells have separated from the tip of the archenteron. (L) Lumen of archenteron has expanded and two coelomic pouches have formed in the developing gut. (M) Immature bipinnaria stage larva. The gut has differentiated into the oesophagus, stomach, and intestine, and coelomic pouches are well-developed. Numerous mesenchymal cells are present. (N) Lateral view of bipinnaria larva. A well-developed, tripartite gut is present, as well as the hydropore canal, which connects one of the coelomic pouches to the exterior of the animal. (O) Bipinnaria larvae with tripartite gut and two elongate coelomic pouches. Green algae are present in the stomach of the animal, showing that it has started feeding. Abbreviations are as follows: GV, Germinal vesicle; PB, polar bodies; FM, Fertilization membrane; VP, Vegetal Plate; A, Archenteron; MeCs, Mesenchymal Cells; CP, Coelomic pouch; St, Stomach; Oe, Oesophagus, HG, Hind gut. Scale bars are 75μm.

In the early bipinnaria larva (Fig 1N), the developing oesophagus bends towards the oral surface and fuses with a pronounced depression, the oral cavity. The lower intestine bends orally and forms the anus. The anus is located on the post-oral lobe (Fig 1N), one of two prominent lobes on the oral surface of the larva. The other lobe, the pre-oral lobe, overhangs the mouth and forms the oral hood (Fig 1N). These lobes are encircled by the pre-oral and post-oral ciliary bands, dense accumulations of ciliated cells. The pre-oral band encircles the anterior end of the larva and the oral hood, while the post-oral ciliary band surrounds the posterior portion of the larva and connects to the lower lip of the oral cavity (Fig 1O, Fig 3A).

### Bipinnaria larvae have two distinct populations of muscle cells

To investigate the molecular basis of the development of larval structures and identify cell types, molecular markers were used to characterize tissues and cells in *A. rubens* larvae at different developmental time points. Musculature was visualized using an antibody against Myosin Heavy Chain (Fig 2A), which has been shown to label fore-gut muscles in sea urchin larvae (Annunziata et al. 2014). In bipinnaria, MHC-immunoreactivity is present in two populations of cells: longitudinal muscles encircling the oesophagus (Fig 2C-D) and bilateral clusters of 6-8 cells near the aboral surface (Fig 2B, E-F). Oesophageal muscle cells are elongate with a prominent nucleus but do not individually encircle the oesophagus (Fig 2D). These cells produce a strong peristaltic contraction, moving food particles through the digestive system (Supp Fig 4A-B). Immunoreactivity is strongest in cells forming the oesophageal-stomach sphincter (Fig 2C). Aboral muscle cells have similar elongate morphologies to oesophageal muscles (Fig 2E-F) but form a clearly distinct structure capable of producing contractions in the aboral surface (Supp Fig 4C).

**Figure 2.**
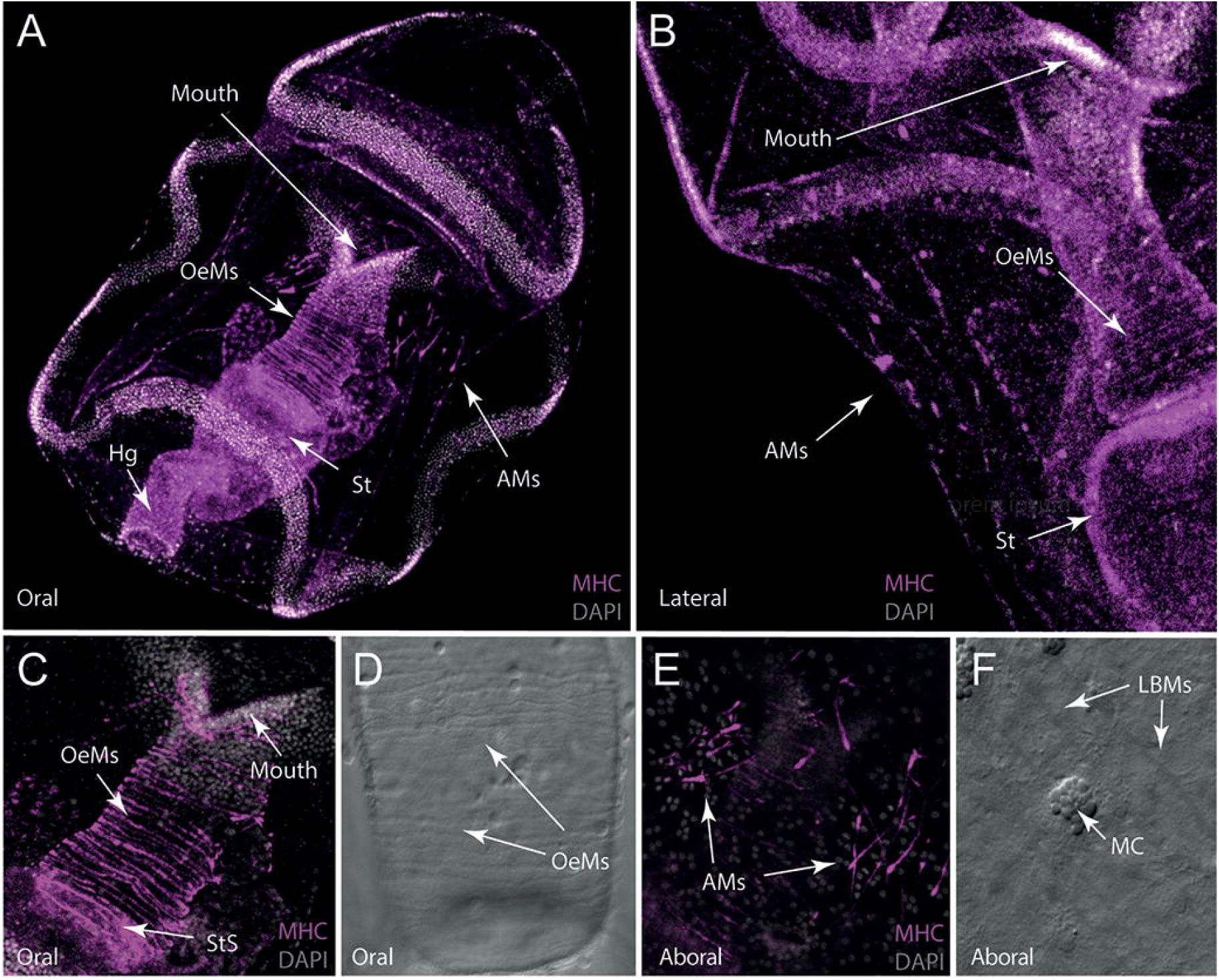
Musculature of bipinnaria larva of *A. rubens*. (A) False colour image showing the oral view of Myosin Heavy Chain (MHC) immunoreactivity and cell nuclei visualized with DAPI (grey). Strong MHC immunostaining can be seen in three populations of muscle cells: fibrous longitudinal muscles encircling the oesophagus, and two clusters of aboral muscles associated with the dorsal surface. (B) Lateral view showing a high magnification view of MHC-immunoreactive aboral muscles located on the dorsal surface of the larva, as well as the longitudinally arranged, fibrous, oesophageal muscles. (C) Oral view showing a high magnification view of MHC-immunoreactive longitudinal fibrous muscles surrounding the oesophagus. (D) Bright-field (with DIC) image showing the fibrous oesophageal muscles surrounding the stomach, which exhibit MHC-immunoreactivity as shown in (C). (E) High magnification image showing MHC-immunoreactivity in aboral muscle cells. These cells are arranged into two clusters on the dorsal surface of the animal and are thicker at one end with a tapering morphology at the other end. (F) Bright-field (with DIC) image of aboral muscle cells and their proximity to mucous cells embedded in the ectoderm of the larva. Abbreviations are as follows: OeMs, oesophageal muscles; AMs, aboral muscles; St, stomach, Sts, Stomach sphincter; MC, mucous cell; Hg, hind gut.

**Figure 3.**
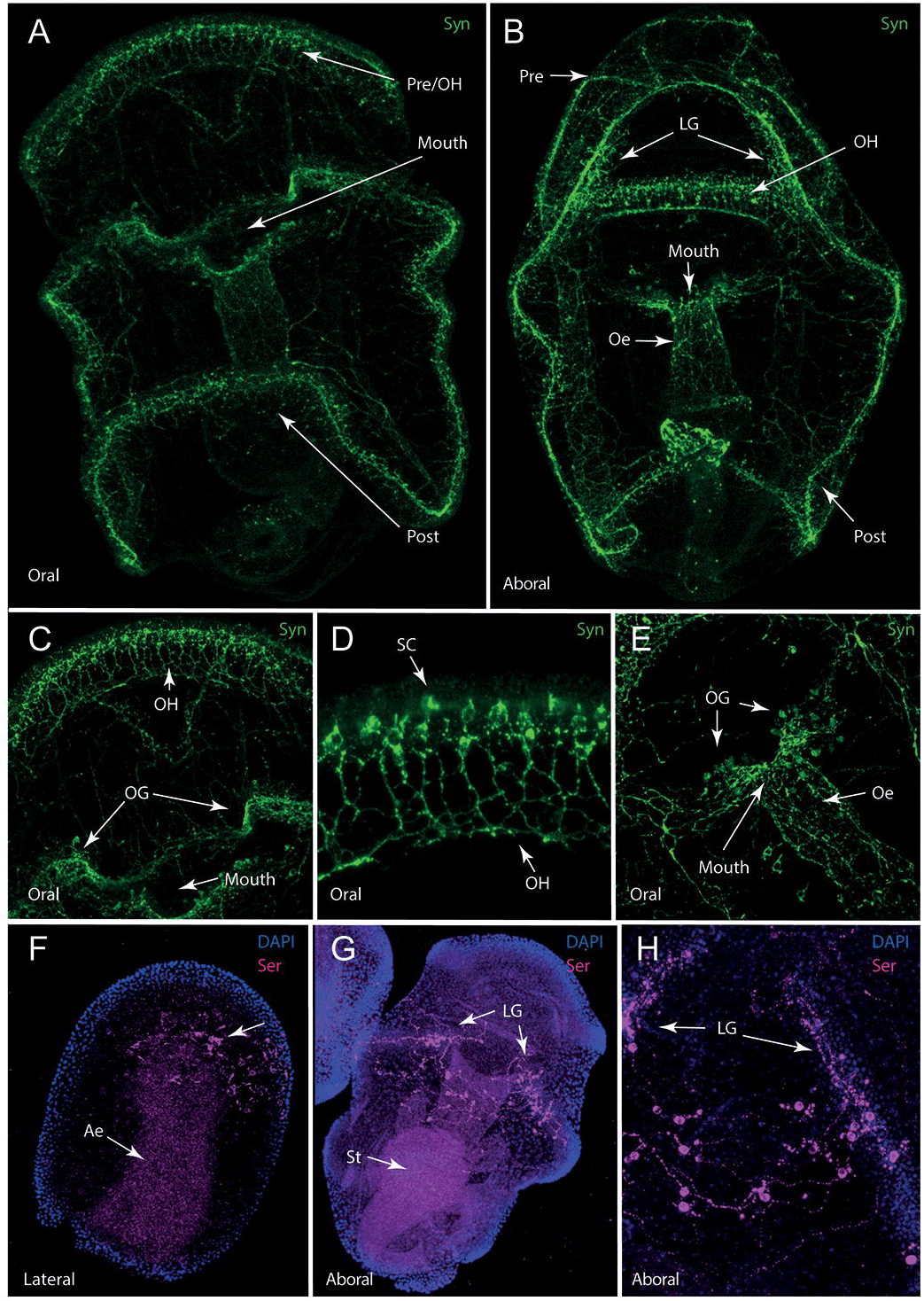
Architecture and morphology of the nervous system in *A. rubens* embryos and larvae. (A) Oral view showing synaptotagmin-immunoreactivity revealed by 1E11 antibodies in the *A. rubens* bipinnaria larva. Strong immunoreactivity is present in the ciliary bands, around the mouth, and along the gut. In addition to these major areas of labelling, synaptotagmin-immunoreactive projections are also present in other regions of the larva. (B) Synaptotagmin-immunoreactivity in a ventral view of a bipinnaria larva. Strong staining is present in the pre-oral ciliary band and the oral hood, as well as in the lateral ganglia and post-oral ciliary band. Innervation can be seen lining the oesophagus, as well as around the mouth. (C) High magnification image of synaptotagmin-immunoreactive neurons in the oral hood with orally-directed projections. Additionally, labelling of clustered neurons of the oral ganglia can be seen on either side of the mouth. (D) High magnification image of synaptotagmin-immunoreactive neurons of the oral hood shown in (C), which can be interpreted as sensory cells with downward projecting processes. (E) High magnification image of synaptotagmin-immunoreactive neuronal cell bodies of the oral ganglia proximal to the mouth and with immunostained processes distributed along the length of the oesophagus. (F) Gastrula stage embryo stained with serotonin antibodies and with cell nuclei labeled using DAPI. Immunoreactivity is strongest in numerous ectodermal cells, interpreted to be neuronal precursors. Immunostaining in the archenteron is interpreted as background staining. (G) Ventral view of an immature bipinnaria showing serotonin immunoreactive cells and processes along the ventral surface. These cells are interpreted to be serotonergic neurons, and are arranged into two clusters in the lateral ganglia. Immunostaining in the stomach is interpreted as background staining. (F) High magnification image of serotonin immunoreactive cells in the lateral ganglia, showing stained cell bodies and neuronal processes. Abbreviations are as follows: Pre, pre-oral ciliary band; OH, oral hood; Post, post-oral ciliary band; LG, lateral ganglia; Oe, oesophagus; OG, oral ganglia; SC, sensory cells.

### Bipinnaria larvae have an extensive nervous system

We next aimed to characterize the *A. rubens* bipinnaria nervous system and how it innervates both the musculature and remainder of the larva. The full extent of the larval nervous system was visualised using the pan-neuronal marker 1E11, which labels the synaptic vesicle trafficking protein synaptotagmin (Fig 3A-B) (Burke et al. 2006). Synaptotagmin immunoreactivity in cells of the larval nervous system is concentrated in the ciliary bands, oral hood and mouth, with projections forming a meshwork around the oesophagus and extending loosely across the rest of the body (Fig 3A-E). Each ciliary band is innervated with a prominent central fibre surrounded by synaptotagmin-immunoreactive neuronal cells laterally positioned on each side (Supp Fig 5A). This central fibre seems to be formed from a bundle of nerve fibres and is least organised where it passes over the oral surface of the post-oral lobe (Fig 3A). Numerous neurites of variable length project laterally from the ciliary band, primarily towards the oral surface (Supp Fig 5B).

A population of neurons in the oral hood have short apical processes (Fig 3D) and may correspond with a well-characterized population of sensory neurons in the apical organ of other echinoderms (Byrne et al. 2007). Synaptotagmin-immunoreactive neurites and neuronal projections extend orally from the basal surface of these cells towards the oral pit, forming a prominent band around its anterior (Fig 3C-D). On either side of the mouth are two prominent ganglia, from which projections connect to both the post-oral ciliary band nerve and form a network surrounding the oesophagus (Fig3 C, E). Gut innervation is concentrated around the oesophagus with limited immunoreactivity in the mid- and hindgut, although some synaptotagmin-immunoreactive cells were identified around the anus (Fig 3A). Outside of the ciliary bands and feeding apparatus, the only neuronal cell bodies observed were a loose cluster of cells spanning the mid-aboral surface (Supp figure 6).

### Neuronal complexity of A. rubens bipinnaria larvae

In other echinoderm larvae, multiple neuronal types have been identified using markers for neurotransmitters, neuropeptides or specific transcription factors (Burke et al. 2006; Slota and McClay 2018; Wood et al. 2018). We have used several antibodies raised against these markers to characterize the bipinnaria nervous system of *A. rubens*. In echinoderms, the neurotransmitter serotonin is present in a subset of apical organ neurons, anterior ganglia considered to be the central nervous system of larvae (Byrne et al. 2007). We used an anti-serotonin antibody to investigate the development and extent of the serotonergic nervous system in *A. rubens* (Fig 3F-H, Supp Fig 6). Serotonin-immunoreactive cells were first observed at the gastrula stage (~46hpf) as a scattered cluster on one side of the animal half of the embryo (Fig 3F, Supp Fig 7). In one-week-old larvae, serotonergic neurons form two clusters at the anterior end of the developing post-oral ciliary band. These develop into the lateral ganglia on the aboral surface of two-week-old larvae (Fig 3G, Supp Fig 7C). These ganglia are connected across the aboral surface by a loose network of cells and projections (Fig 3H). We found no evidence of serotonin-immunoreactivity on the oral surface or in the oral region at any stage analysed.

Previous studies have identified sub-populations of neurons that express genes encoding different neuropeptide precursors in sea urchin larvae (Wood et al. 2018). To further characterize neuronal cell types in the ciliary band nervous system of *A. rubens*, we used antibodies raised against the RNA-binding protein ELAV (Garner et al. 2016), considered a specific neuronal marker, a sea urchin AN-peptide-type neuropeptide (Perillo et al. 2018) and the *A. rubens* SLFMaide neuropeptide (S2). ELAV-immunoreactivity was observed in cell bodies of some, but not all ciliary band neurons of two-week-old larvae (compare Fig3A and 4A). Immunoreactivity was also observed in the lower gut and stomach (Fig 4 A-B). Conversely, AN-peptide antibodies label four sets of 4-5 pyramidal cells in the lateral portions of both ciliary bands. The cells of each set are connected by lateral neuronal projections in the ciliary bands (Fig 4C-E). Two sets of cells are paired on either side of the pre-oral ciliary band running oral-aborally (Fig 4C) and the remaining cells are paired on either side of the post-oral ciliary band beside the post-oral lobe (Fig 4D-E). Antibodies against S2 label a small bilaterally arranged set of ciliary band neurons in the anterior loop of the post-oral ciliary band (Fig 4F-G). Short S2-immunoreactive projections extend posteriorly from the ciliary bands across the aboral surface. Importantly, these neurons are located anteriorly to the lateral ganglia and appear distinct from the adjacent serotonin-immunoreactive cells. Collectively, these data suggest a degree of sub-functionalisation within the ciliary band.

**Figure 4.**
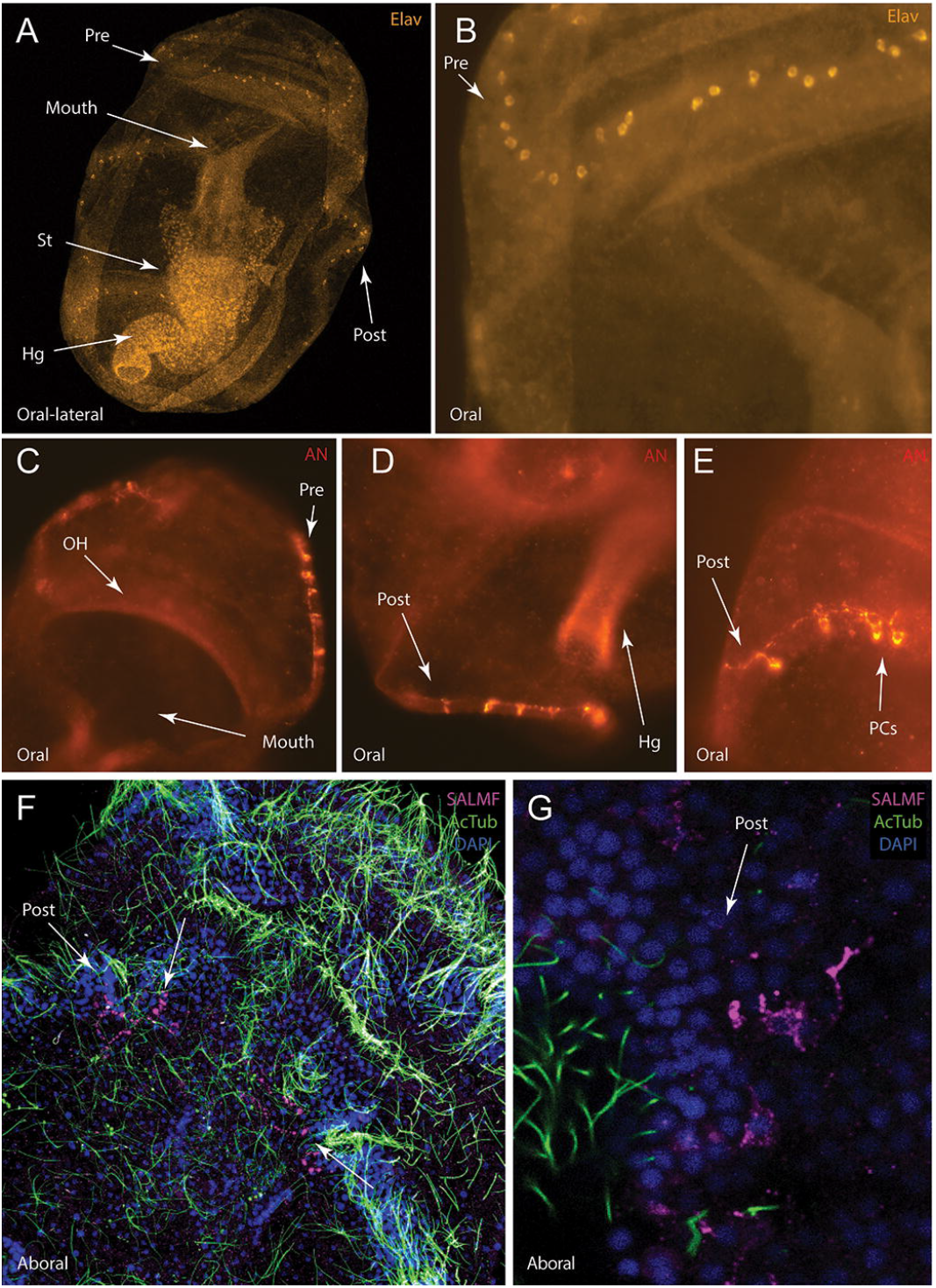
Neuronal subpopulations expressing ELAV or neuropeptides (AN-peptide, SALMFamide S2) in *A. rubens* larvae. (A) Oral view of the *A. rubens* bipinnaria showing ELAV-immunoreactivity. Immunostaining is strongest in distinct subpopulations of cells within the pre-oral and post-oral ciliary bands. (B) High magnification image of ELAV-immunoreactive cells in the pre-oral ciliary band and oral hood. Note that neuronal processes are not labeled by the ELAV-antibodies and immunoreactivity is present in some, but not all, ciliary band neurons. (C) High magnification image of the oral hood and pre-oral ciliary band showing AN-peptide immunoreactive neuronal cells and processes. (D) High magnification image of the post-oral ciliary band showing AN-peptide immunoreactive neuronal cells and processes. (E) High magnification image of the post-oral ciliary band showing AN-peptide immunoreactive neuronal cells and processes. (F) S2-immunoreactive processes (pink) in the anterior loop of the post-oral ciliary band and in projections extending onto the oral surface in a larva that has also been labelled for acetylated tubulin (green) and with a nuclear marker (DAPI; blue). (G) High magnification image showing S2-immunoreactive cells and their processes in the post-oral ciliary band. Abbreviations are as follows: Pre; pre-oral ciliary band; Post, post-oral ciliary band; St, stomach; Hg, hindgut; OH, oral hood; PCs, Pyramidal cells.

Antibodies against the *A. rubens* pedal peptide-type neuropeptide ArPPLN1b (Lin et al. 2017) and SoxB2, a transcription factor expressed during nervous system development (Cunningham et al. 2008), labelled a sub-population of neurons in the oral region of the bipinnaria. An ArPPLN1b-immunoreactive cluster of cells with associated projections surrounds the mouth and anterior end of the oesophagus (Fig 5B-C) and resemble synaptotagmin-immunoreactive oral neurons.

**Figure 5.**
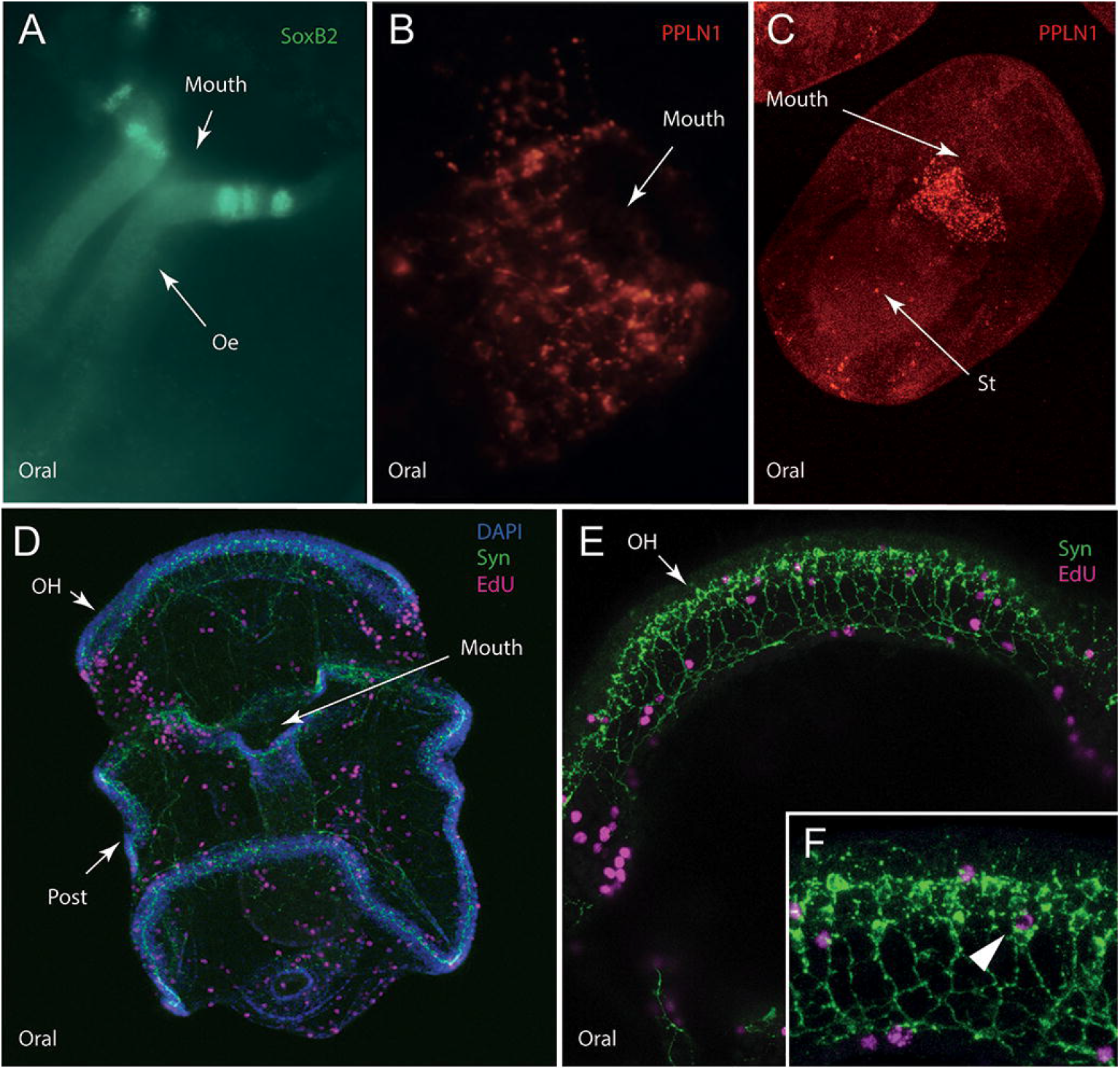
Neuronal subpopulations expressing SoxB2 or the neuropeptide ArPPLN1b and cells undergoing proliferation in *A. rubens* larvae. (A) Immunostaining of a sub-population of cells expressing the transcription factor SoxB2 in mouth region of a bipinnaria larva. (B) Antibodies to the neuropeptide ArPPLN1b reveal immunostained cells and processes. around the mouth of a bipinnaria larva. (C) As in bipinnaria larvae, in immature bipinnaria ArPPLN1b -immunoreactivity is strongest in a subpopulation of cells around the mouth. (D) Cell proliferation in a bipinnaria larva labelled using the marker EdU together with immunostaining for synaptotagmin using 1E11 antibodies. EdU-positive cells are located throughout the larva, but are most strongly concentrated in the ciliary bands. Cell proliferation is most evident along the edges of the pre-oral lobe and along the edges of the mouth. (E) Enlargement of D showing the oral hood (arrow). (F) Two slide projection of a part of the oral hood showing an EdU-positive cell that is co-stained with 1E11 antibodies is present. Abbreviations are as follows: Oe, oesophagus; St, stomach; OH, oral hood; Post, post-oral ciliary band.

ArPPLN1b-immunoreactive cells are first observed around the newly formed oesophagus of one-week-old larvae (Supp Fig 8) as a loose network of cells and become prominently localised in projections encircling the entrance to the oesophagus by two weeks post fertilization (Fig 5C). Conversely SoxB2 was observed only in three small, discrete clusters of cells around the mouth, just above the entrance to the oesophagus (Fig 5A). These cells are morphologically distinct from ArPPLN1b-immunoreactive cells, consistent with different roles for the molecular markers; i.e. SoxB2 is involved in neuronal cell precursor specification whereas ArPPLN1b is likely functional in differentiated neurons (Cunningham et al. 2008).

In summary, these data reveal a remarkable diversity of neuronal cell types, highlighting the complexity of the larval nervous system in *A. rubens*.

### Cell proliferation in larval growth and development

As larval growth proceeds, the bipinnaria nervous system of *A. rubens* increases in complexity and the ciliary bands become more defined. Therefore, we investigated the role of cell proliferation in development of the larval nervous system, with particular focus on the growth of the ciliary bands and associated innervation.

To identify sites of cell proliferation during growth, we used 5-ethynyl-2’-deoxuridine (EdU), a nucleoside analogue of thymidine that is incorporated into newly synthesized DNA and thus marks dividing cells (Zeng et al. 2010). In the blastula and early gastrula, there were no specific patterns of cell proliferation, with cells dividing throughout the embryo (Supp Fig 9). Two-week-old bipinnaria show continued cell proliferation at reduced levels throughout the embryo with cell division concentrated in the ciliary bands (Fig 5D-E). Within the ciliary bands, proliferation is most apparent on either side of the pre-oral lobe and the edges of the oral pit (Fig 5D). Cell proliferation is also prominent in the coelomic pouches.

To better understand where cell proliferation in the ciliary band occurred relative to the tissues of the nervous system, EdU marked samples were co-stained with 1E11. Generally, co-localisation was absent, consistent with the notion that terminally differentiated neurons do not divide (Jukam and Desplan 2010). However, a small subset of synaptotagmin-immunoreactive cells appear to show nuclear EdU (Fig 5F), which we interpret as dividing differentiated neurons.

## Discussion

### Conservation of the bipinnaria body plan

Our findings (Fig 1) reveal that the general body plan and early developmental progression of the bipinnaria larvae of *A. rubens* is similar to that of previously studied asteroids (Hinman et al. 2003; Nakajima et al. 2004; Dyachuk and Odintsova 2013; Pernet et al. 2017). This conservation of development and structure is remarkable given the considerable phylogenetic divergence between studied asteroid species (Supp Fig 10). *P. miniata* and *A. rubens* have nearly identical bipinnaria larvae despite representing different asteroid super-orders that diverged ~200 million years ago (Lafay et al. 1995; Linchangco et al. 2017). The only notable variation between species is the speed of larval development. The feeding bipinnaria is present by 3 dpf in Patiria miniata and Patiria pectinifera (Murabe et al. 2021) but not until 5-6 dpf in Pisaster ochraceus and *A. rubens* (Pia et al. 2012). Our work also shows more rapid developmental rates in larvae reared at higher temperatures (Supp Fig 1), supporting the view that temperature, not inherent developmental regulation, is the main factor determining speed of larval development (Cataldo et al. 2005).

In indirect developing asteroids, morphological deviations from the bipinnaria body plan are only apparent at relatively late stages in larval development, such as in members of the genus Luidia (Wilson 1978), or in post-metamorphic juveniles (Morris 2009; Pernet et al. 2017). Such limited variation in early bipinnaria development suggests a strong evolutionary or developmental constraint on this stage for species utilizing this larval strategy. Our work, and the ubiquity of the bipinnaria across asteroid phylogeny, further support the hypothesis that an indirect developing, feeding and free-swimming larvae represents the ancestral mode of development for sea stars (Raff and Byrne 2006). We also note that despite the diversity of non-bipinnaria asteroid larvae, none retain bipinnaria derived structures and there appear to be no intermediate forms (McEdward and Janies 1997). We therefore hypothesize the presence of a developmental switch between two stable larval strategies (planktotrophic bipinnaria and largely direct developing lecithotrophs) rather than gradual morphological shifts.

### Neuronal complexity in the early larvae

Our results show that just two weeks post-fertilization, the bipinnaria has developed a complex nervous system extending throughout the body with distinct central and peripheral components. The central nervous system consists of the concentration of neuronal cell bodies in the anterior apical region, while the peripheral nervous system comprises the neurons along the ciliary bands and around the mouth as well as projections innervating the rest of the body (Fig 6). This complexity is consistent with previous studies of the bipinnaria nervous system (Nakajima et al. 2004) and subdivisions into central and peripheral components are a shared organisational feature of the nervous systems of echinoids and asteroids (Hinman and Burke 2018). This study is the first, however, to show the scale of neuronal diversity in the early bipinnaria. Within the centralized nervous system, we have identified at least two centres of neuronal complexity: the anterior end of the post-oral ciliary band around the apical organ and the cells of the oral opening. Additional neuronal diversity is present along the remainder of the ciliary bands (Fig 6). Since the primary role of planktonic larvae is to find and consume food, neuronal complexity within structures involved in feeding and locomotion suggests that the majority of these neurons play a role in the coordination of these processes.

**Figure 6.**
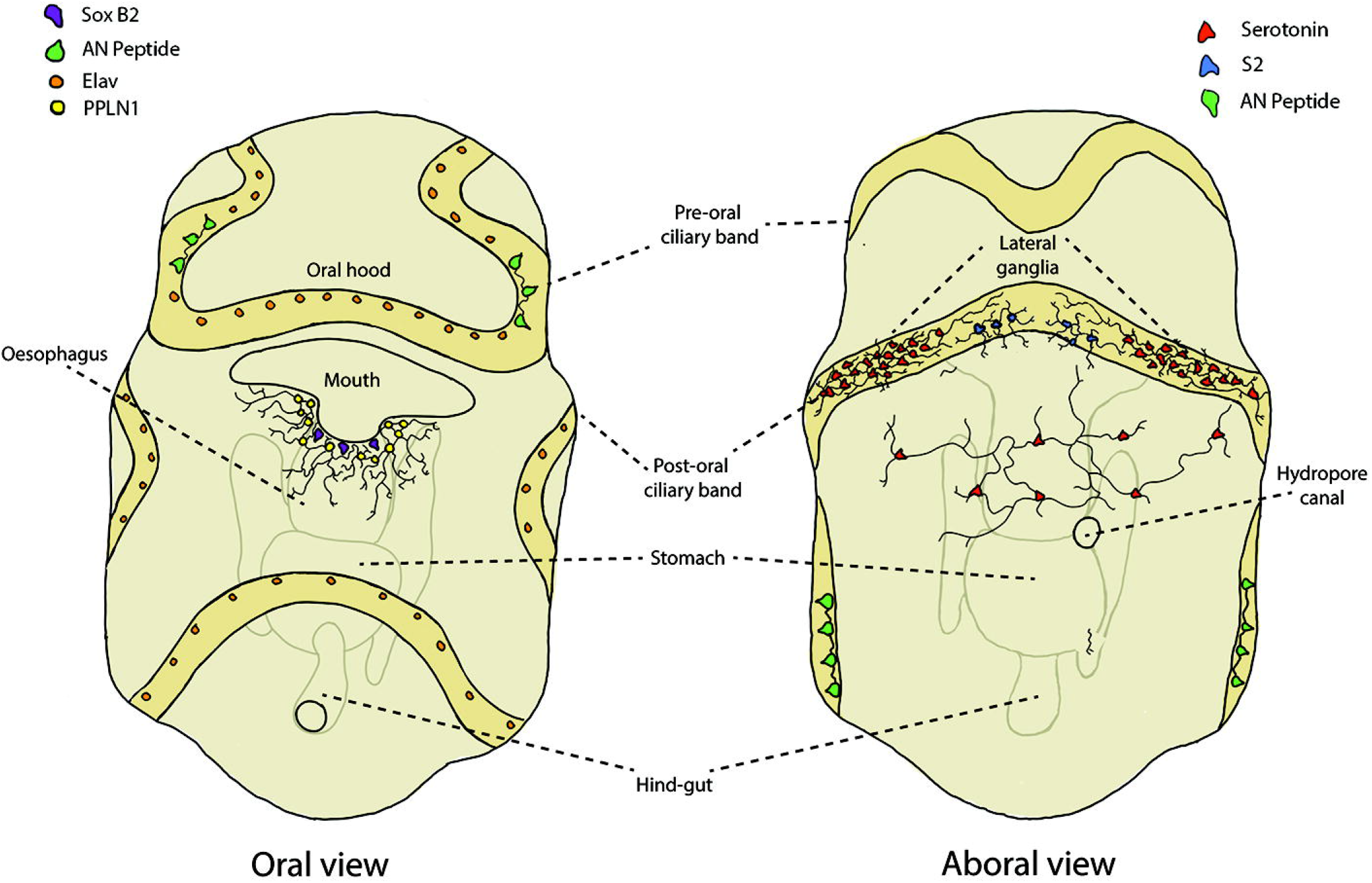
Diagrammatic representation of neuronal subpopulations in *A. rubens* bipinnaria larva. (Left) Oral view showing subpopulations of neurons expressing ELAV, SoxB2 and the neuropeptides AN-peptide and ArPPLN1b. The mouth appears to be a center of neuronal complexity, with at least two distinct subpopulations of neurons present. (Right) Aboral view showing subpopulations of serotonergic neurons and neurons expressing neuropeptides (S2 and AN peptide).

In the oral region, our results show that the neuropeptide ArPPLN1b is expressed in a loose network of neurons surrounding the anterior portion of the oesophagus (Fig 5B). While the expression pattern of this neuropeptide has not been described previously in larvae, studies in adult *A. rubens* have revealed that it causes muscle relaxation and so is also known as starfish myorelaxant peptide (Kim et al. 2016; Lin et al. 2017). The proximity of ArPPLN1b-immunoreactivity to the larval oesophageal muscles suggests a potentially conserved role in neuromuscular signalling between larvae and adults, despite the full breakdown and reassembly of the nervous system during metamorphosis. Detailed neuroanatomical work will, however, be required to determine if interactions exist between these neurons and oesophageal muscles.

We also identified expression of the neurogenic transcription factor SoxB2 in the oral region, distinct from ArPPLN1b-immunoreactive neurons around the opening of the oesophagus (Fig 5A). SoxB2 is associated with the nervous system throughout the animal kingdom (Royo et al. 2011) and is specifically involved in neurogenesis in the echinoid pleuteus larvae foregut, where it is the first marker expressed in neuronal precursors (Anishchenko et al. 2018). Oral neurons develop in-situ in echinoid larvae (Wei et al. 2011) and we hypothesize the presence of a similar system in asteroids, with SoxB2 expressed in neuronal precursors and ArPPLN1b expressed in differentiated oral/oesophageal neurons.

The RNA binding protein ELAV is expressed in a subset of cells along the length of both ciliary bands, in agreement with previous studies of echinoid and asteroid larvae (Yankura et al. 2013; Garner et al. 2016). Unlike these studies, we found no evidence of ELAV-immunoreactive cells in the oral region despite the extensive neuronal array labelled by synaptotagmin antibodies. This suggests a limitation of ELAV as a consistent marker of all major neuronal fields, at least in *A. rubens* larvae.

AN-peptide-immunoreactivity is localised in a unique pattern of two paired clusters of cells in the pre- and post-oral ciliary bands (Figs 4C-E, 8A), distinct from all other neuronal markers in this study. This pattern is also distinct from the expression pattern observed in larvae of *S. purpuratus* (Perillo et al. 2018), where AN peptide antibodies label lateral ganglia and apical organ neurons that co-express serotonin. That the expression pattern of this neuropeptide is so different between asteroids and echinoids suggests that it may have evolved distinct functions in each class. Therefore, further investigation of AN peptide expression and function in echinoderm larvae may provide valuable insights into the neuronal complexity and evolutionary history of early bipinnaria larvae.

### The serotonergic nervous system and the apical organ

The serotonergic nervous system has been extensively studied across the larvae of the Echinodermata, although research has primarily focused on echinoids and asteroids (Moss et al. 1994; Chee and Byrne 1999; Nakajima et al. 2004; Jarvela et al. 2016). The development and localisation of serotonergic neurons varies considerably between echinoderm classes, but is considered in all groups to mark the neurons of the apical organ (Byrne et al. 2007). In *A. rubens* we first observed serotonin-immunoreactive cells at gastrulation, with immunoreactivity concentrated in the lateral ganglia by two-weeks post-fertilisation. A subset of serotonin-immunoreactive cells connect the lateral ganglia across the aboral surface. As the only non-ciliary neurons on the aboral surface, these cells may also play a role in coordinating contractions of the aboral muscles that allow the bipinnaria to reverse (Strathmann 1971). As with ArPPLN1b, detailed neuroanatomical work will be required to confirm an interaction between the two cell types. Based on analysis of the expression of transcripts encoding the precursor of the neuropeptide S2 (F-type SALMFamide precursor) using in-situ hybridisation (Mayorova et al. 2016), it has been suggested that this neuropeptide type is a marker of the apical organ in asteroids. Our immunohistochemical analysis of S2 expression indicates that it is localised in cells at the anterior end of the post-oral band but that these neurons are located slightly anteriorly to the main concentrations of serotonergic cells in the lateral ganglia (Fig 6 right) and may represent a distinct population of cells. Further work is required to determine the function of these cells in the bipinnaria larvae.

Among studied asteroids, the extent of the serotonergic nervous system in *A. rubens* is most similar to Pisaster ochraceus (Moss et al. 1994), but greatly restricted when compared to Patiriella regularis, where serotonin immunoreactive cells also extend into the oral region and around the oral hood (Byrne et al. 2007). Because *P. regularis* belongs to the order Valvatida while *P. ochraceus* and *A. rubens* belong to the order Forcipulatida (Supp Fig 10), this may reflect a shared ancestral loss of this part of the serotonergic nervous system, or elaboration in *P. regularis* during the ~200 million years since their evolutionary divergence.

## Conclusion

Our work herein sheds light on the complexity of neuronal cell types and musculature present in the seemingly simple *A. rubens* larvae. This work provides the foundations for further characterisation of larval development in *A. rubens* in conjunction with new genomic and transcriptomic resources for this species. Furthermore, while we have continued to expanded knowledge of asteroid development across evolutionary timescales, future work investigating the development of bipinnaria and non-bipinnaria larvae will be necessary to provide a more complete understanding of larval evolution in the Asteroidea.

## Acknowledgments

The author would like to thank Wendy Hart for help in animal husbandry and lab experiments, Dr Natalie Wood for precious advices on immunostaining, the UCL imaging facility and other members of the Oliveri lab: Kirn Doal and Emanuele Astoricchio.

## References

Anishchenko E, Arnone MI, D’Aniello S. 2018. SoxB2 in sea urchin development: implications in neurogenesis, ciliogenesis and skeletal patterning. EvoDevo 9:5.

Annunziata R, Perillo M, Andrikou C, Cole AG, Martinez P, Arnone MI. 2014. Pattern and process during sea urchin gut morphogenesis: The regulatory landscape. genesis 52:251–68.

Boveri T. 1893. An Organism Produced Sexually without Characteristics of the Mother. The American Naturalist 27:222–32.

Budd GC. 2008. Asterias rubens: Common Starfish. In: Tyler-Walters H, Hiscock K, editors. Marine Life Information Network: Biology and Sensitivity Key Information Reviews,[on-line] Plymouth: Marine Association of the United Kingdom.

Burke RD. 1983. The structure of the larval nervous system of Pisaster ochraceus (Echinodermata: Asteroidea). Journal of Morphology 178:23–35.

Burke RD, Osborne L, Wang D, Murabe N, Yaguchi S, Nakajima Y. 2006. Neuron-specific expression of a synaptotagmin gene in the sea urchin Strongylocentrotus purpuratus. Journal of Comparative Neurology 496:244–51.

Byrne M, Cisternas P. 2002. Development and distribution of the peptidergic system in larval and adult Patiriella: Comparison of sea star bilateral and radial nervous systems. Journal of Comparative Neurology 451:101–14.

Byrne M, Koop D, Strbenac D, Cisternas P, Balogh R, Yang JYH, Davidson PL, Wray G. 2020. Transcriptomic analysis of sea star development through metamorphosis to the highly derived pentameral body plan with a focus on neural transcription factors. DNA Research 27.

Byrne M, Nakajima Y, Chee FC, Burke RD. 2007. Apical organs in echinoderm larvae: insights into larval evolution in the Ambulacraria. Evolution & Development 9:432–45.

Byrne M, Selvakumaraswamy P. 2002. Phylum Echinodermata: Ophiuroidea. In: Young C, editor. Atlas of Marine Invertebrate Larvae London: Academic Press. p. 483–98.

Cai W, Kim C-H, Go H-J, Egertová M, Zampronio CG, Jones AM, Park NG, Elphick MR. 2018. Biochemical, Anatomical, and Pharmacological Characterization of Calcitonin-Type Neuropeptides in Starfish: Discovery of an Ancient Role as Muscle Relaxants. Front Neurosci 12:382.

Cary GA, Cameron RA, Hinman VF. 2018. EchinoBase: Tools for Echinoderm Genome Analyses. In: Kollmar M, editor. Eukaryotic Genomic Databases: Methods and Protocols. Methods in Molecular Biology New York, NY: Springer. p. 349–69.

Cary GA, Hinman VF. 2017. Echinoderm development and evolution in the post-genomic era. Developmental Biology 427:203–11.

Cary GA, McCauley BS, Zueva O, Pattinato J, Longabaugh W, Hinman VF. 2020. Systematic comparison of sea urchin and sea star developmental gene regulatory networks explains how novelty is incorporated in early development. Nature Communications 11:6235.

Cataldo D, Boltovskoy D, Hermosa JL, Canzi C. 2005. Temperature-Dependent Rates Of Larval Development In Limnoperna Fortunei (Bivalvia: Mytilidae). Journal of Molluscan Studies 71:41–46.

Chee F, Byrne M. 1999. Development of the Larval Serotonergic Nervous System in the Sea Star Patiriella regularis as Revealed by Confocal Imaging. The Biological Bulletin 197:123–31.

Cunningham DD, Meng Z, Fritzsch B, Casey ES. 2008. Cloning and developmental expression of the soxB2 genes, sox14 and sox21, during Xenopus laevis embryogenesis. Int J Dev Biol 52:999–1004.

Davidson EH, Erwin DH. 2006. Gene Regulatory Networks and the Evolution of Animal Body Plans. Science 311:796–800.

Davidson EH, Rast JP, Oliveri P, Ransick A, Calestani C, Yuh C-H, Minokawa T, Amore G, Hinman V, Arenas-Mena C, Otim O, Brown CT, Livi CB, Lee PY, Revilla R, Rust AG, Pan Z jun, Schilstra MJ, Clarke PJC, Arnone MI, Rowen L, Cameron RA, McClay DR, Hood L, Bolouri H. 2002. A Genomic Regulatory Network for Development. Science 295:1669–78.

Driesch H. 1892. The potency of the first two cleavage cells in echinoderm development. Experimental production of partial and double formations. Foundations of experimental embryology.

Dyachuk V, Odintsova N. 2013. Larval myogenesis in Echinodermata: conserved features and morphological diversity between class-specific larval forms of Echinoidae, Asteroidea, and Holothuroidea. Evolution & Development 15:5–17.

Dylus DV, Czarkwiani A, Stångberg J, Ortega-Martinez O, Dupont S, Oliveri P. 2016. Large-scale gene expression study in the ophiuroid Amphiura filiformis provides insights into evolution of gene regulatory networks. EvoDevo 7:2.

Elphick MR, Newman SJ, Thorndyke MC. 1995. Distribution and action of SALMFamide neuropeptides in the starfish Asterias rubens. Journal of Experimental Biology 198:2519–25.

Garner S, Zysk I, Byrne G, Kramer M, Moller D, Taylor V, Burke RD. 2016. Neurogenesis in sea urchin embryos and the diversity of deuterostome neurogenic mechanisms. Development 143:286–97.

Gemmill JF. 1914. VII. The development and certain points in the adult structure of the starfish asterias rubens, L. Philosophical Transactions of the Royal Society of London Series B, Containing Papers of a Biological Character 205:213–94.

Haesaerts D, Jangoux M, Flammang P. 2005. The attachment complex of brachiolaria larvae of the sea star Asterias rubens (Echinodermata): an ultrastructural and immunocytochemical study. Zoomorphology 124:67–78.

Hall MR, Kocot KM, Baughman KW, Fernandez-Valverde SL, Gauthier MEA, Hatleberg WL, Krishnan A, McDougall C, Motti CA, Shoguchi E, Wang T, Xiang X, Zhao M, Bose U, Shinzato C, Hisata K, Fujie M, Kanda M, Cummins SF, Satoh N, Degnan SM, Degnan BM. 2017. The crown-of-thorns starfish genome as a guide for biocontrol of this coral reef pest. Nature 544:231–34.

Hart MW, Byrne M, Smith MJ. 1997. Molecular Phylogenetic Analysis of Life-History Evolution in Asterinid Starfish. Evolution 51:1848–61.

Hinman VF, Burke RD. 2018. Embryonic neurogenesis in echinoderms. WIREs Developmental Review 7.

Hinman VF, Nguyen AT, Cameron RA, Davidson EH. 2003. Developmental gene regulatory network architecture across 500 million years of echinoderm evolution. PNAS 100:13356–61.

Jarvela AMC, Hinman V. 2014. A Method for Microinjection of Patiria minata Zygotes. JoVE (Journal of Visualized Experiments) e51913.

Jarvela AMC, Yankura KA, Hinman VF. 2016. A gene regulatory network for apical organ neurogenesis and its spatial control in sea star embryos. Development 143:4214–23.

Jukam D, Desplan C. 2010. Binary fate decisions in differentiating neurons. Current Opinion in Neurobiology, Development 20:6–13.

Kim C-H, Kim EJ, Go H-J, Oh HY, Lin M, Elphick MR, Park NG. 2016. Identification of a novel starfish neuropeptide that acts as a muscle relaxant. J Neurochem 137:33–45.

Lacalli TC, Gilmour THJ, West JE. 1990. Ciliary band innervation in the bipinnaria larva of Pisaster ochraceus. Philosophical Transactions of the Royal Society of London Series B: Biological Sciences 330:371–90.

Lafay B, Smith AB, Christen R. 1995. A Combined Morphological and Molecular Approach to the Phylogeny of Asteroids (Asteroidea: Echinodermata). Systematic Biology 44:190–208.

Lin M, Egertová M, Zampronio CG, Jones AM, Elphick MR. 2017. Pedal peptide/orcokinin-type neuropeptide signaling in a deuterostome: The anatomy and pharmacology of starfish myorelaxant peptide in Asterias rubens. Journal of Comparative Neurology 525:3890–3917.

Linchangco GV, Foltz DW, Reid R, Williams J, Nodzak C, Kerr AM, Miller AK, Hunter R, Wilson NG, Nielsen WJ, Mah CL, Rouse GW, Wray GA, Janies DA. 2017. The phylogeny of extant starfish (Asteroidea: Echinodermata) including Xyloplax, based on comparative transcriptomics. Molecular Phylogenetics and Evolution 115:161–70.

Mah C, Foltz D. 2011. Molecular phylogeny of the Forcipulatacea (Asteroidea: Echinodermata): systematics and biogeography. Zoological Journal of the Linnean Society 162:646–60.

Mayorova TD, Tian S, Cai W, Semmens DC, Odekunle EA, Zandawala M, Badi Y, Rowe ML, Egertová M, Elphick MR. 2016. Localization of Neuropeptide Gene Expression in Larvae of an Echinoderm, the Starfish Asterias rubens. Front Neurosci 10.

McEdward LR. 1995. Evolution of pelagic direct development in the starfish Pteraster tesselatus (Asteroidea: Velatida). Biological Journal of the Linnean Society 54:299–327.

McEdward LR, Janies DA. 1997. Relationships among development, ecology, and morphology in the evolution of Echinoderm larvae and life cycles. Biological Journal of the Linnean Society 60:381–400.

McEdward LR, Miner BG. 2001. Larval and life-cycle patterns in echinoderms. Canadian Journal of Zoology.

Morris VB. 2009. On the sites of secondary podia formation in a juvenile echinoid: growth of the body types in echinoderms. Dev Genes Evol 12.

Moss C, Burke RD, Thorndyke MC. 1994. Immunocytochemical localization of the neuropeptide S1 and serotonin in larvae of the starfish Pisaster ochraceus and Asterias rubens. Journal of the Marine Biological Association of the United Kingdom 74:61–71.

Murabe N, Okumura E, Chiba K, Hosoda E, Ikegami S, Kishimoto T. 2021. The Starfish Asterina pectinifera: Collection and Maintenance of Adults and Rearing and Metamorphosis of Larvae. In: Carroll DJ, Stricker SA, editors. Developmental Biology of the Sea Urchin and Other Marine Invertebrates: Methods and Protocols. Methods in Molecular Biology New York, NY: Springer US. p. 49–68.

Nakajima Y, Kaneko H, Murray G, Burke RD. 2004. Divergent patterns of neural development in larval echinoids and asteroids. Evolution & Development 6:95–104.

Newman SJ, Elphick MR, Thorndyke MC. 1995. Tissue distribution of the SALMFamide neuropeptides S1 and S2 in the starfish Asterias rubens using novel monoclonal and polyclonal antibodies. I. Nervous and locomotory systems. Proceedings of the Royal Society of London Series B: Biological Sciences 261:139–45.

Odekunle EA, Semmens DC, Martynyuk N, Tinoco AB, Garewal AK, Patel RR, Blowes LM, Zandawala M, Delroisse J, Slade SE, Scrivens JH, Egertová M, Elphick MR. 2019. Ancient role of vasopressin/oxytocin-type neuropeptides as regulators of feeding revealed in an echinoderm. BMC Biol 17:60.

Perillo M, Paganos P, Mattiello T, Cocurullo M, Oliveri P, Arnone MI. 2018. New Neuronal Subtypes With a “Pre-Pancreatic” Signature in the Sea Urchin Stongylocentrotus purpuratus. Front Endocrinol 9.

Pernet B, Livingston BT, Sojka C, Lizárraga D. 2017. Embryogenesis and larval development of the seastar Astropecten armatus. Invertebrate Biology 136:121–33.

Peter IS, Davidson EH. 2015. Genomic Control Process Elsevier.

Peterson KJ, Arenas-Mena C, Davidson EH. 2000. The A/P axis in echinoderm ontogeny and evolution: evidence from fossils and molecules. Evolution & Development 2:93–101.

Pia TS, Johnson T, George SB. 2012. Salinity-induced morphological changes in Pisaster ochraceus (Echinodermata: Asteroidea) larvae. Journal of Plankton Research 34:590–601.

Raff RA. 2008. Origins of the other metazoan body plans: the evolution of larval forms. Philosophical Transactions of the Royal Society B: Biological Sciences 363:1473–79.

Raff RA, Byrne M. 2006. The active evolutionary lives of echinoderm larvae. Heredity 97:244–52.

Royo JL, Maeso I, Irimia M, Gao F, Peter IS, Lopes CS, D’Aniello S, Casares F, Davidson EH, Garcia-Fernández J, Gómez-Skarmeta JL. 2011. Transphyletic conservation of developmental regulatory state in animal evolution. PNAS 108:14186–91.

Ruiz-Ramos DV, Schiebelhut LM, Hoff KJ, Wares JP, Dawson MN. 2020. An initial comparative genomic autopsy of wasting disease in sea stars. Molecular Ecology 29:1087–1102.

Scheltema RS. 1986. Long-distance dispersal by planktonic larvae of shoal-water benthic invertebrates among central Pacific islands. Bulletin of Marine Science 39:241–56.

Semmens DC, Mirabeau O, Moghul I, Pancholi MR, Wurm Y, Elphick MR. 2016. Transcriptomic identification of starfish neuropeptide precursors yields new insights into neuropeptide evolution. Open Biology 6:150224.

Slota LA, McClay DR. 2018. Identification of neural transcription factors required for the differentiation of three neuronal subtypes in the sea urchin embryo. Developmental Biology 435:138–49.

Stewart MJ, Stewart P, Rivera-Posada J. 2015. De novo assembly of the transcriptome of Acanthaster planci testes. Mol Ecol Resour 15:953–66.

Strathmann RR. 1971. The feeding behavior of planktotrophic echinoderm larvae: Mechanisms, regulation, and rates of suspensionfeeding. Journal of Experimental Marine Biology and Ecology 6:109–60.

Thompson JR, Paganos P, Benvenuto G, Arnone MI, Oliveri P. 2020. Post-metamorphic skeletal growth in the sea urchin Paracentrotus lividus and implications for body plan evolution. bioRxiv 2020.10.09.332957.

Tian S, Egertová M, Elphick MR. 2017. Functional Characterization of Paralogous Gonadotropin-Releasing Hormone-Type and Corazonin-Type Neuropeptides in an Echinoderm. Front Endocrinol (Lausanne) 8:259.

Wei Z, Angerer RC, Angerer LM. 2011. Direct development of neurons within foregut endoderm of sea urchin embryos. PNAS 108:9143–47.

Wilson DP. 1978. Some Observations on Bipinnariae and Juveniles of the Starfish Genus Luidia. Journal of the Marine Biological Association of the United Kingdom 58:467–78.

Wood NJ, Mattiello T, Rowe ML, Ward L, Perillo M, Arnone MI, Elphick MR, Oliveri P. 2018. Neuropeptidergic Systems in Pluteus Larvae of the Sea Urchin Strongylocentrotus purpuratus: Neurochemical Complexity in a “Simple” Nervous System. Front Endocrinol 9.

Wray GA. 1992. The evolution of larval morphology during the post-Paleozoic radiation of echinoids. Paleobiology 18:258–87.

Yankura KA, Koechlein CS, Cryan AF, Cheatle A, Hinman VF. 2013. Gene regulatory network for neurogenesis in a sea star embryo connects broad neural specification and localized patterning. PNAS 110:8591–96.

Young C, Sewell M, Rice M (Eds.). 2002. Atlas of Marine Invertebrate Larvae London: Academic Press.

Zamora LN, Delorme NJ, Byrne M, Sewell MA. 2020. Lipid and protein utilization during lecithotrophic development in the asteroid Stegnaster inflatus, with a review of larval provisioning in lecithotrophic echinoderms. Marine Ecology Progress Series 641:123–34.

Zeng C, Pan F, Jones LA, Lim MM, Griffin EA, Sheline YI, Mintun MA, Holtzman DM, Mach RH. 2010. Evaluation of 5-ethynyl-2′-deoxyuridine staining as a sensitive and reliable method for studying cell proliferation in the adult nervous system. Brain Research 1319:21–32.

Zhang Y, Yañez Guerra LA, Egertová M, Zampronio CG, Jones AM, Elphick MR. 2020. Molecular and functional characterization of somatostatin-type signalling in a deuterostome invertebrate. Open Biol 10:200172.

