## Supplemental Information for "The development and neuronal complexity of bipinnaria larvae of the sea star *Asterias rubens*"

#### Extended Materials and Methods

*Collection of adults and husbandry*
Adult *A. rubens* were obtained from a fisherman based at Whitstable in Kent (UK). Animals were initially maintained in large seawater tanks at Queen Mary University of London and prior to embryological work transferred in artificial sea water (ASW) to tanks at University College London. Specimens were sexed by gonad biopsy and males and females isolated in separate tanks. Each aquarium tank contained circulating ASW at ~12°C with a salinity of 34-36ppt, corresponding to the natural habitat of this species (Budd 2008). Adults were fed periodically with the blue mussel *Mytilus edulis*.

*Gamete collection and embryo culture*
For gamete collection and the initiation of embryo cultures we used a procedure modified from Jarvela and Hinman (2014). Specifically, gonads were extracted from one arm of a gravid male and female for each round of fertilization through a small incision in the aboral surface of each arm at the point of fusion with the central disc. Testes were collected dry and kept refrigerated (4˚C) for several days and then gently squeezed to release spermatozoa when needed. Ovaries were kept in ASW (33.5ppt) and shredded with forceps to separate oocytes from the associated follicle cells and remaining ovarian tissue. Oocytes were passed through a 180μm filter to remove large debris, then washed twice in ASW to remove small fragments of remaining gonadic tissue. 1-methyladenine (1-MA) to a final concentration of 10 μM was added to the oocytes to induce germinal vesicle breakdown and oocyte maturation. Oocytes were incubated for 40 minutes (min) at 12˚C and maturation was monitored under the microscope. Post-maturation, eggs were washed in ASW to remove excess 1-MA. 0.2 μl/ml undiluted spermatozoa were added to the eggs and gently mixed. Fertilisation was allowed to proceed for roughly 15 min and was monitored under the microscope before excess spermatozoa were removed and fertilised oocytes re-suspended at a concentration of 300-500 embryos/ml in ASW containing streptomycin (50μg/mL) and penicillin (20U/mL).

Fertilized eggs were incubated at either 12° or 15°C in large petri dishes for optimal gas exchange and embryonic development was allowed to proceed. ASW was replaced every three days and once larvae became free swimming, a small air pump was used to generate a water current in the culture. Once the oral cavity had fully opened, larvae were fed a mixture of *Dunaliella sp* and *Rhodomonas sp* every three days. Developmental progression was checked hourly until embryos had undergone their fifth cleavage, then at least once daily until the culture reached the bipinnaria stage. At each time point, several individuals (>30) were mounted on a glass slide and were visualised using a Zeiss AxioImager M1. Embryos and larvae were collected at different stages for further processing.

*Immunostaining*

For each developmental time point of interest, 15 ml of culture (4000-7000 embryos) was placed on ice until all embryos settled. Excess ASW was removed and concentrated embryos were fixed in 4% paraformaldehyde (PFA:ASW) for 15 min on ice. Samples were permeabilized for either 1 min in methanol or 10 min in phosphate buffered saline containing 1:1000 Tween and 1:10000 Triton X (PBST+Triton) depending on the primary antibody (Supplemental Table 1). Samples treated with methanol were subsequently washed 5 times in PBST and all samples were stored for 24-72 hours in blocking buffer (BB: 1:25 sheep or goat serum and 1:100 bovine serum albumin (BSA) in PBST). Around 100 embryos were processed for each combination of antibodies. BB was removed and samples were incubated with primary antibodies in BB at 37° C for 1.5 h (concentrations given in Supplemental Table 1). Embryos were washed 4 times in PBST and re-suspended in 1:1000 secondary antibody in PBST in the dark for 1 hour at room temperature (RT). Embryos were washed 2 times in PBST and incubated in DAPI (final concentration 1:100000) in PBST for 5 min at RT in the dark. Embryos were finally washed 2 times in PBST and stored in the dark at 4°C until imaging.

*Assays of cell proliferation*

Developing larvae were stained with 5-ethynyl-2'-deoxyuridine (EdU) to visualise dividing cell nuclei using the Click-iT® EdU Alexa Flour® 555HCS kit (Life Technologies). Samples were initially incubated in 1:1000 EdU for two hours and then fixed in 1:25 paraformaldehyde in PBS for 15 min. Each sample was then washed 3 times in PBST and then permeabilized in PBST+Triton for 45 min. Samples were washed a further 2 times in PBST and then permeabilized in methanol for 2 min before a further two washes in PBS and two washes in PBST. Samples were then incubated in BB overnight and immunostained as described above. Following post-secondary antibody PBST washes, samples were incubated in 50 μl of Click-iT® Reaction cocktail for 30 min following the manufacturer's instructions. Samples were then washed with 50 μl of Click-iT® reaction Rinse Buffer for 30 min, before two washes in PBST, and addition of DAPI at a final concentration of 1:100000.

*Light and confocal microscopy*

For differential interference contrast (DIC) images as well as epi-fluorescent images a Zeiss AxioImager M1 microscope was used together with a Zeiss AxioCamHRc camera. Confocal microscopy was carried out at the UCL Imaging Facility using an LSM 800 confocal microscope or Leica SPEinv inverted confocal microscope with sequential scanning. Z-stack images or single slices were taken using confocal microscopy, employing the optimal step size with a pinhole size of 1 au. Images were taken using a

10x lens or 25x lens with immersion oil. Following microscopy, image processing and analysis was carried out using ImageJ version 2.0.0.0.

#### Supplementary Tables

| Primary Antibody | Secondary Antibody | Concentration of Primary Ab | Methanol (Y/N) |
|---|---|---|---|
| Anti-synaptotagmin B (1E11) | Goat anti-Mouse IgG (H+L) Cross-Adsorbed Secondary Antibody, Alexa Fluor 488, | 1:10 | Y |
| Anti-serotonin | Goat anti-Rabbit IgG (H+L) Cross-Adsorbed Secondary Antibody, Alexa Fluor 555, | 1:500 | Y |
| Anti-MHC | Goat anti-Rabbit IgG (H+L) Cross-Adsorbed Secondary Antibody, Alexa Fluor 555, | 1:500 | Y |
| Anti-acetylated-tubulin | Goat anti-Mouse IgG (H+L) Cross-Adsorbed Secondary Antibody, Alexa Fluor 488, | 1:400 | Y |
| Anti-Elav | Goat anti-Rabbit IgG (H+L) Cross-Adsorbed Secondary Antibody, Alexa Fluor 555, | 1:400 | Y |
| Anti-SoxB2 | Goat anti-Rat IgG (H+L) Cross-Adsorbed Secondary Antibody, Alexa Fluor 488, | 1:400 | Y |
| Anti-ArPPLN1b | Goat anti-Rabbit IgG (H+L) Cross-Adsorbed Secondary Antibody, Alexa Fluor 555, | 1:50000 | Y |
| Anti-SLFMide (S2) | Goat anti-Rabbit IgG (H+L) Cross-Adsorbed Secondary Antibody, Alexa Fluor 555, | 1:500 | Y |
| Anti-AN-peptide | Goat anti-Rabbit IgG (H+L) Cross-Adsorbed Secondary Antibody, Alexa Fluor 555, | 1:500 | Y |

**Supplementary Table 1**: Primary and secondary antibody (ab) combinations used in immunostaining experiments to visualise the nervous system and musculature of the bipinnaria larvae of *Asterias rubens*. The final concentration of primary antibodies in blocking buffer is given as well as the name of each secondary antibody. Additionally, we annotated whether or not the antibody combination worked with permeabilisation using methanol.

### Extended results

#### Extended description of development in *A. rubens*

Ooyctes are held in late prophase in the female gonads prior to spawining, as shown by the large germinal vesicle that disappears following 1-MA induced maturation (Fig 1A, B). Mature eggs were on average 160 μm in diameter. The fertilization membrane elevated within minutes of sperm addition but varied in proximity to the surface of the egg. In each batch, development proceeded synchronously with first cleavage following extrusion of the polar bodies (Fig 1C) complete by 3 hpf at 12°C and 2 hpf at 15°C. Subsequent cleavages are holoblastic and largely equal for roughly 9 to 11 cycles occurring on average every 3 h at 12°C and 2 h at 15°C. A large blastocoel is visible in the early blastula corresponding to the ~254 cell stage (Fig 1F). Growth is constrained by the fertilisation membrane until ~24 hpf (Fig 1G), with loss of the membrane accompanied by a pronounced elongation along the animal/vegetal axis and a thickening of the cells in the posterior (vegetal) half of the embryo (Fig 1H). In some embryos, multiple invaginations of the epithelium gave the blastula a wrinkled appearance (Supp Fig 2), similar to the phenomenon previously noted in some primarily polar, lecithotrophic asteroid species (Henry et al. 1991). However, the presence of these invaginations were not consistent between batches and had no apparent impact on later development. As expected, embryos developed consistently more rapidly at higher temperatures (Supp Fig 1).

By 30 hpf at 15°C (36hpf for 12 C), the vegetal plate starts to flatten and thicken into a denser mass of cells as a precursor to gastrulation (Fig 1I), producing an invaginating archenteron that reaches half the length of the embryo by 46 hpf. During gastrulation, the lumen of the archenteron gradually decreases in diameter as the walls become noticeably thicker and the archenteron extends in length (Fig1J and K). By 70 hpf, mesenchyme cells arising from the tip of the archenteron are apparent in the blastocoel (Fig1K) and these bud-off into the blastocoel throughout gastrulation (Fig 1K-M). By 92 hpf two differently sized coelomic pouches are apparent and the central part of the archenteron expands to form the proto-stomach (Fig 1L). The immature bipinnaria (122 hpf) has a tripartite gut with an oesophagus, stomach and hind-gut (Fig 1M), and mesenchymal cells that have ingressed into the blastocoel start to show considerable morphological differentiation (Supp Fig 3).

In the early bipinnaria larva the developing oesophagus bends towards the oral surface and fuses with a pronounced depression known as the oral cavity (Fig 1N). The lower intestine bends orally away from the stomach and fuses with the ectoderm to form the anus. The anus is located on the post-oral lobe (Fig 1N), one of two prominent lobes on the oral surface of the larva. The other lobe, termed the pre-oral lobe, overhangs the mouth and forms the oral hood (Fig1N). These lobes are encircled by the pre- and post-oral ciliary bands, dense accumulations of ciliated cells. The pre-oral band encircles the anterior end of the larva and the oral hood, while the post-oral ciliary band surrounds the posterior portion of the larva and connects to the lower lip of the oral cavity (Fig 1O, Fig 3A).

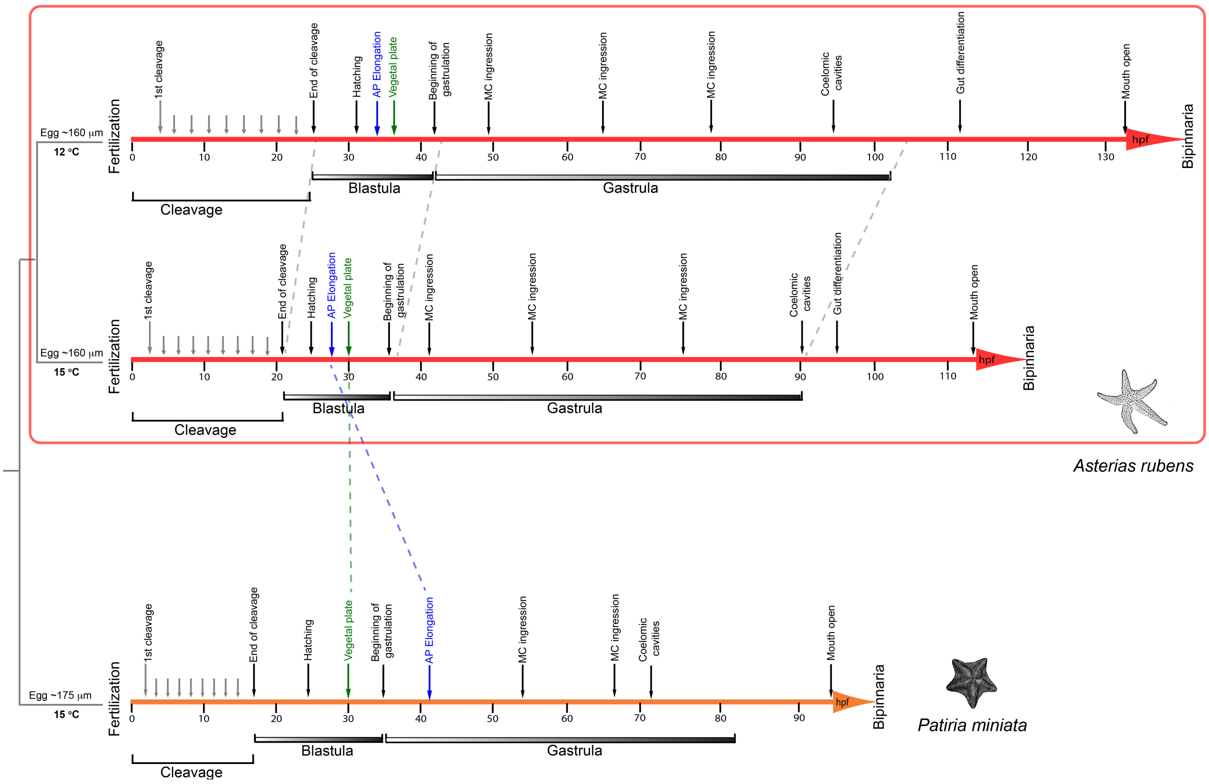


**Supplementary Figure 1.** Comparison of the timing of developmental processes in the bipinnaria larvae of *Asterias rubens* at two different temperatures and in the sea-star *Patiria miniata*. The egg of *P. miniata* has a slightly larger diameter (~175 μm) to that of *A. rubens* (~160 μm). Development in *A. rubens* occurs faster at 15°C than at 12°C, with the larval mouth opening around 113 hpf at 15°C and around 133 hpf at 12°C. All proceeding developmental stages are reached faster at 15°C than at 12°C. Development to the opening of the mouth is more rapid in *P. miniata* than for *A. rubens* at any temperature, occurring around 95 hpf. The rate of development is similar between *A. rubens* larvae at 15°C and *P. miniata* until gastrulation. AP elongation occurs before thickening of the vegetal plate in *A. rubens*, but after thickening of the vegetal plate in *P. miniata.* Abbreviations are as follows: hpf, hours post fertilization.

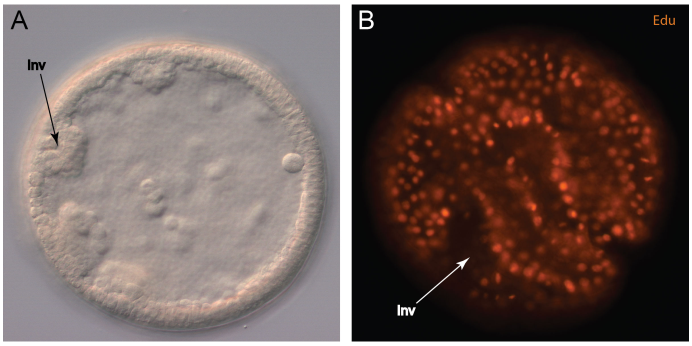


**Supplementary Figure 2**. Wrinkled blastula of *Asterias rubens.* (A) Blastula stage embryo prior to loss of the fertilization membrane showing invaginations of cells into blastocoel. (B) Surface view of EdU-labelled blastula stage embryo showing wrinkled appearance of embryo due to invaginations. Abbreviations are as follows: Inv, invaginations.

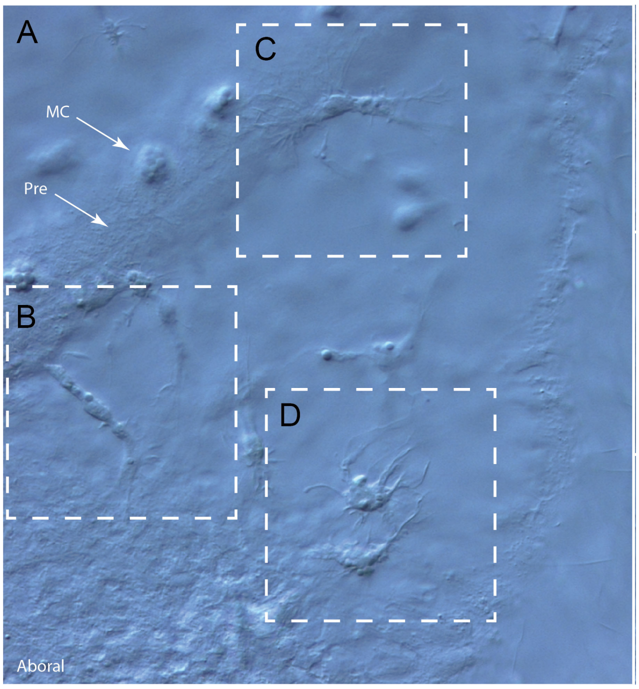


**Supplementary Figure 3**. Diversity of mesenchymal cells in early bipinnaria larvae of *Asterias rubens*. (A) Aboral surface of a 1 week old bipinnaria larvae showing diversity of mesenchymal cells close to a portion of the pre-oral ciliary band and granular mucous cells on the aboral surface. (B) An elongate columnar cell. (C) A neuron-like cell with neurite-like processes associated with the pre-oral ciliary band (D) A phagocyte-like cell with long processes and a granular appearance. Abbreviations are as follows: Pre, pre-oral ciliary band; MC, mucous cell.

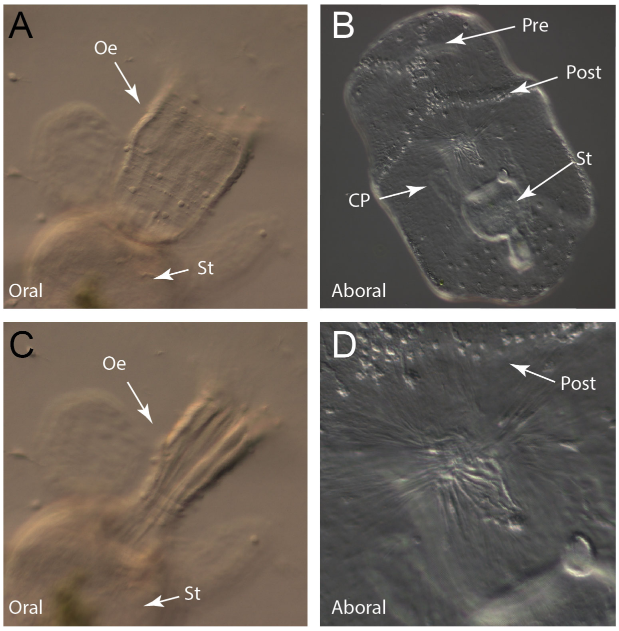


**Supplementary Figure 4.** Contractions of the larval musculature. (A) Oral view of oesophagus with muscles in a relaxed state prior to peristaltic contraction. (B) Aboral view of bipinnaria larvae showing separating ciliary bands and contraction of aboral muscles below the aboral surface. (C) Oral view of oesophagus with muscles undergoing peristaltic contraction. (D) Magnification of contraction of the aboral surface caused by the action of aboral muscles below the surface. Abbreviations are as follows: Oe, oesophagus; St, stomach; Pre, pre-oral ciliary band; Post, post-oral ciliary band; CP, coelomic pouch.

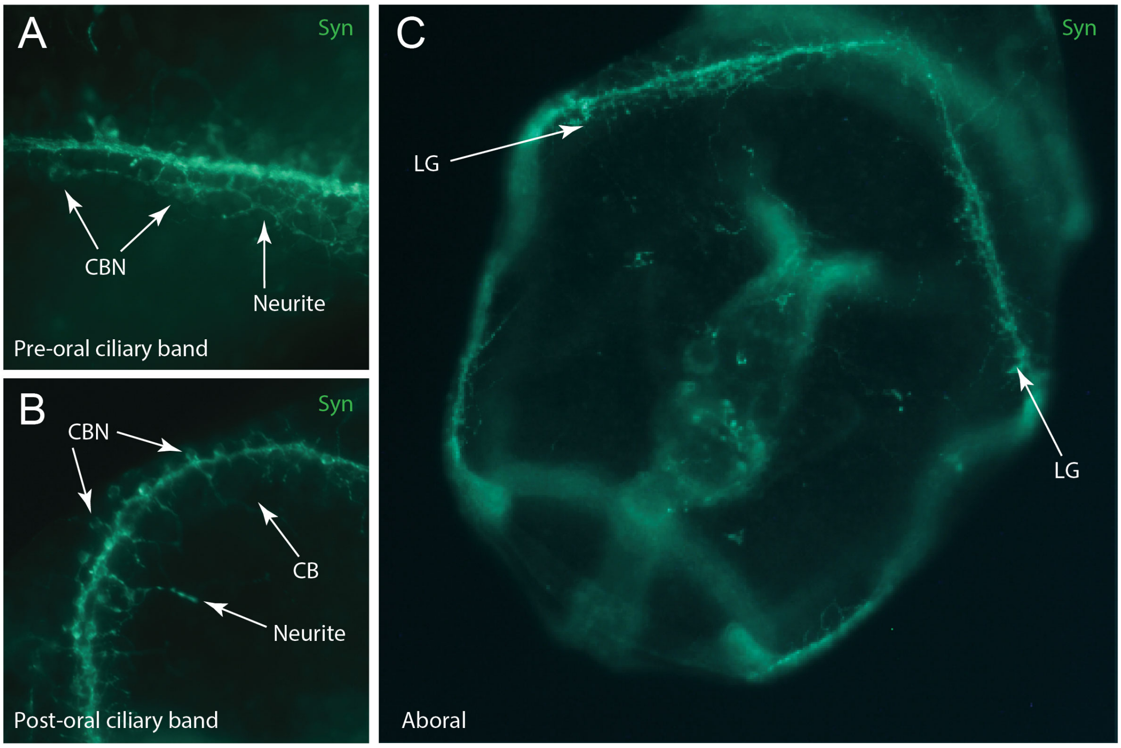


**Supplementary Figure 5.** False colour image of details of ciliary band innervation in the bipinnaria larvae of *Asterias rubens*. Synaptotagmin-immunoreactivity is shown in green. (A) Synaptotagmin-immunoreactivity revealed by 1E11 antibodies in a portion of the pre-oral ciliary band. Nerve fibres are bundled together to form a main ciliary band nerve with neurites forming a loose network extending from the central nerve. (B) Synaptotagmin-immunoreactivity in a portion of the post-oral ciliary band. Neurites and cell bodies are situated laterally alongside the ciliary band nerve with cell bodies roughly evenly spaced. (C) Aboral view of synaptotagmin-immunoreactivity on the aboral surface of a larva showing concentration of immunoreactivity in the lateral ganglia of the post-oral ciliary band and restriction of non-ciliary aboral immunoreactivity to a loose network of cells between the lateral ganglia. Abbreviations are as follows: CBN, Ciliary band neuron; Pre, pre-oral ciliary band; Post, post-oral ciliary band; CB, Cell body; LG, Lateral ganglia.

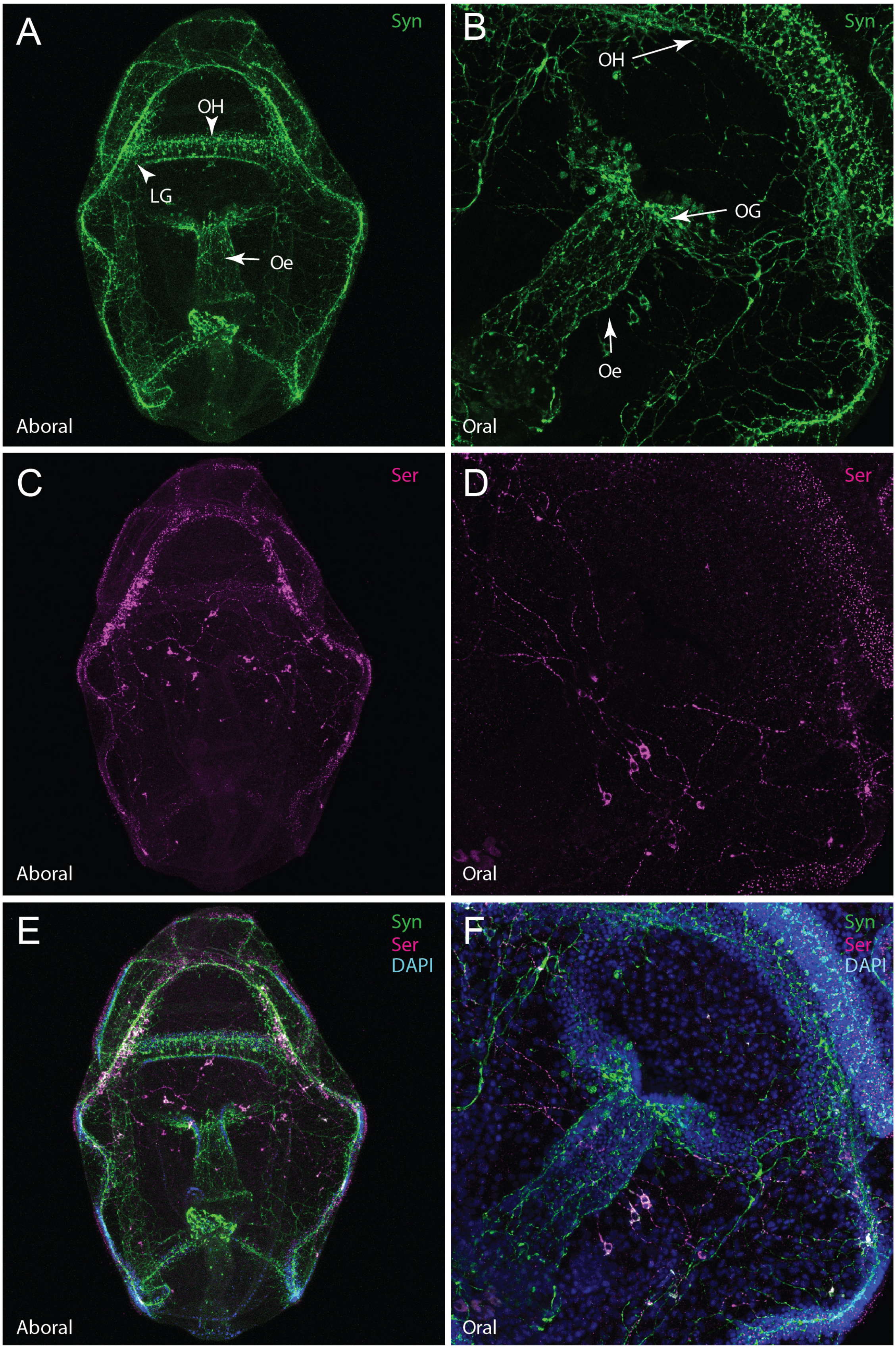


**Supplementary Figure 6.** False colour image of co-localization of synaptotagmin (green) and serotonin (magenta) in the bipinnaria larvae of *Asterias rubens*. (A) Maximum projection aboral view of synaptotagmin-immunoreactivity revealed by 1E11 antibodies in the bipinnaria larvae. Synaptotagmin immunoreactivity is concentrated in the ciliary bands, oral hood and around the oesophagus. (B) Maximum projection detail of oral region in aboral view showing synaptotagmin-immunoreactivity concentrated in the oral ganglia and encircling the oesophagus. (C) Maximum projection aboral view of bipinnaria larvae stained with anti-serotonin antibodies. Serotonin immunoreactivity is concentrated in the lateral ganglia in the post-oral ciliary band and a loose network of cells spanning the aboral surface between the ganglia. (D) Maximum projection detail of oral region in aboral view showing anti-serotonin immunoreactivity concentrated in a small number of cells and projections spanning the aboral surface. (E) Multi-channel combined image of a maximum projection aboral view of a bipinnaria larva co-stained with anti-synaptotagmin (1E11) and anti-serotonin antibodies and with cell nuclei labelled using DAPI (blue). Co-localisation of synaptotagmin and serotonin can be observed in the lateral ganglia (white) and in cells spanning the aboral surface. (F) Multi-channel combined image of oral region in aboral view showing co-localization of synaptotagmin, serotonin and DAPI. Limited co-localization (white) occurs in the cells spanning the oral surface. Abbreviations are as follows: OH, Oral hood; LG, Lateral ganglia; Oe, Oesophagus; OG, Oral ganglia.

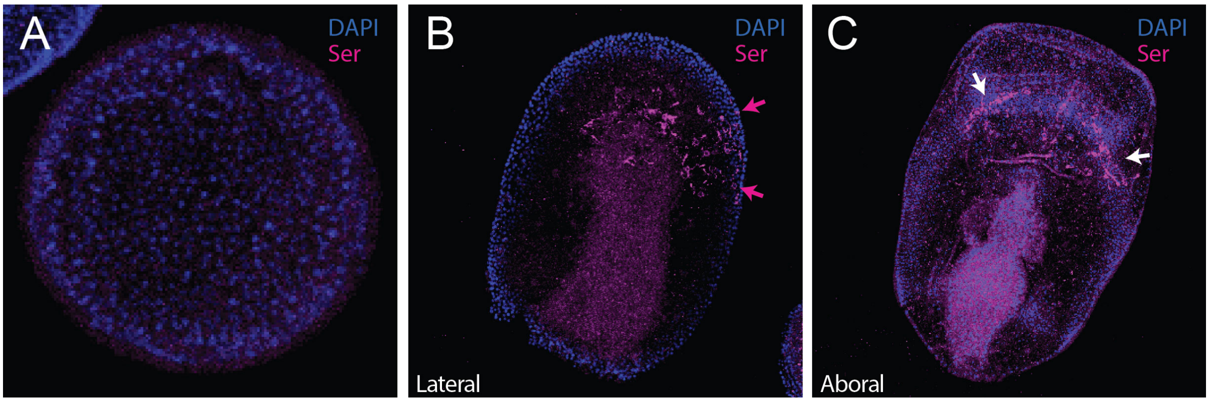


**Supplementary Figure 7.** False colour image of development of the serotonergic nervous system. DAPI is blue and serotonin-immunoreactive cells are in magenta. (A) Blastula stage larva stained with anti-serotonin antibodies and with cell nuclei labelled using DAPI showing absence of serotonin immunoreactivity. (B) Gastrula stage embryo stained with anti-serotonin antibodies and with cell nuclei labelled using DAPI. Immunoreactivity is strongest in numerous ectodermal cells, interpreted to be neuronal precursors. Immunostaining in the archenteron is interpreted as background staining. (C) Ventral view of an immature bipinnaria showing serotonin-immunoreactive cells and processes along the ventral surface. These cells are interpreted to be serotonergic neurons and are arranged into two clusters in the lateral ganglia. Immunostaining in the stomach is interpreted as background staining.

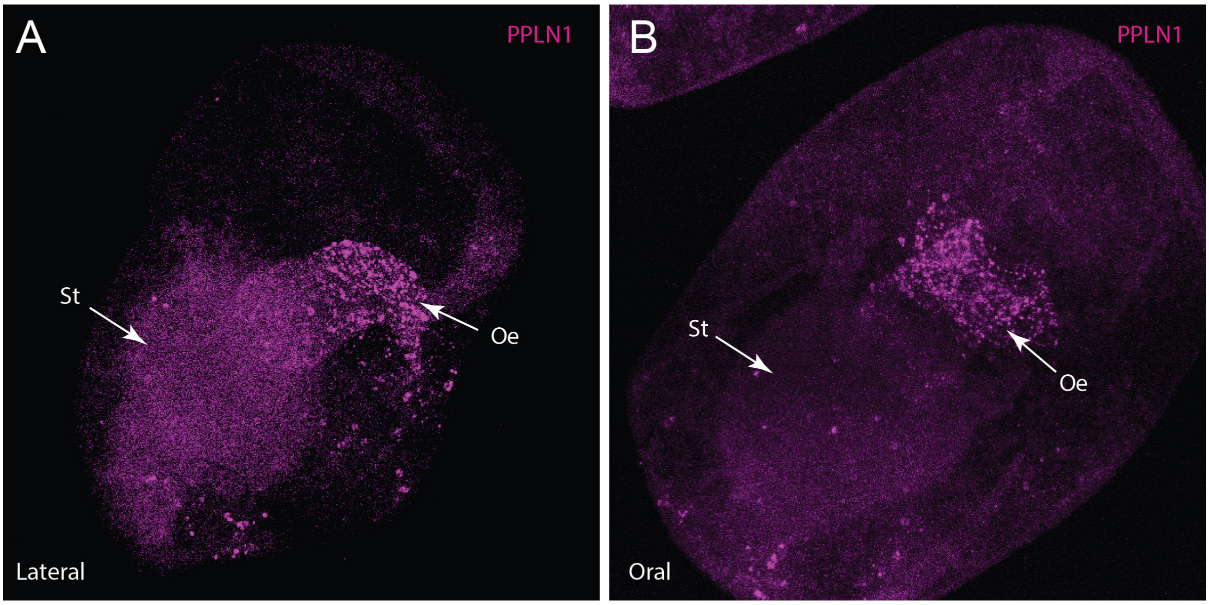


**Supplementary Figure 8.** False colour image of development of the ArPPLN1b localization in the nervous system. (A) In an immature bipinnaria, antibodies against the neuropeptide ArPPLN1b reveal immunostained cells around the developing oesophagus where it bends towards to oral surface. (B) In a more-developed bipinnaria larva, antibodies against the neuropeptide ArPPLN1b reveal immunostained cells and processes around the mouth. Abbreviations are as follows: Oe, Oesophagus; St, stomach.

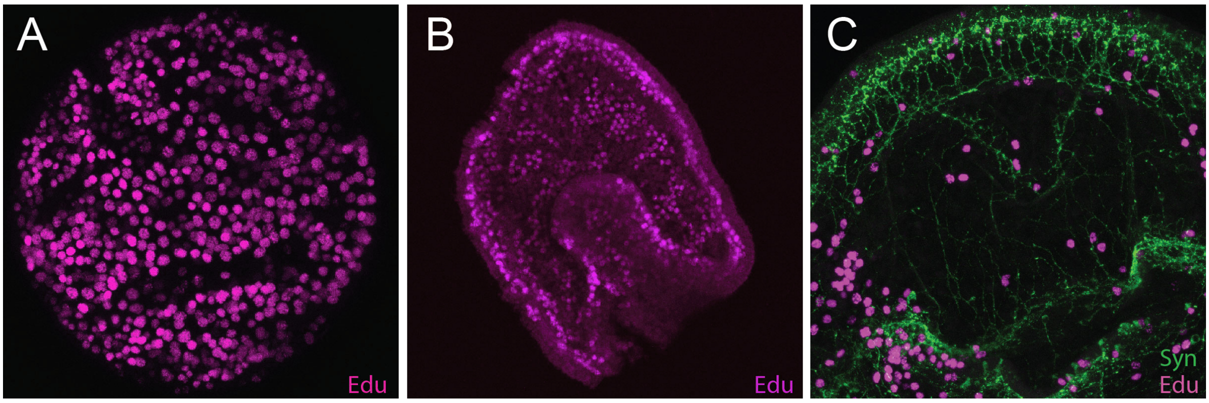


**Supplementary Figure 9.** False colour image of cell proliferation throughout development in the bipinnaria larvae. EdU-stained cells shown in magenta, while synaptotagmin-immunoreactive cells are in green. (A) Blastula stage embryo labelled using the marker EdU. EdU-positive cells are located throughout the embryo. (B) Gastrula stage embryo labelled using the marker EdU. EdU-positive cells are located throughout the embryo. (C) Detail of the oral region of a two-week old bipinnaria larvae labelled using the marker EdU and anti-synaptotagmin antibody (1E11). EdU-positive cells are located throughout the region but are concentrated at the edges of the oral ganglia and ciliary bands.

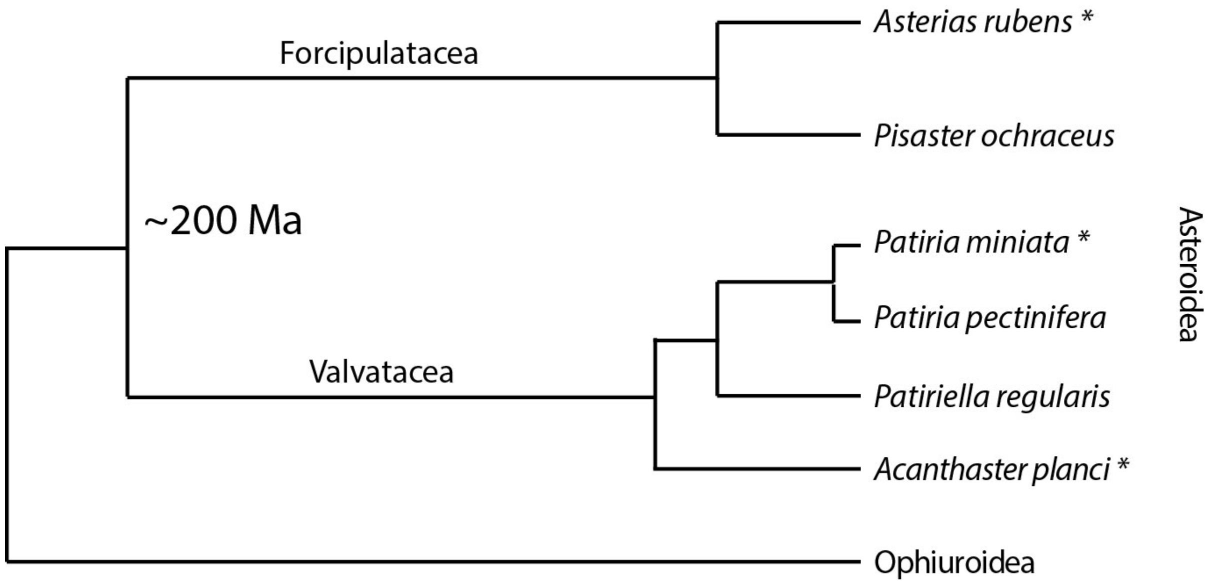


**Supplementary Figure 10.** Phylogenetic representation of relationships between the asteroid species mentioned in the main text. The species studied represent two asteroid super-orders (Forcipulatacea and Valvatacea) that diverged ~200 million years ago. *Asterias rubens* and *Pisaster ochraceus* are Forcipulatid asteroids belonging to the family Asteriidae respectively. The valvatid asteroids *Patiria miniata*, *Patiria pectinifera* and *Patiriella regularis* are all members of the family Asterinidae. The sister group to all asteroids represented is the Ophiuroidea. Asterisks denote species with a sequenced genome.

#### Figure Captions Abbreviations List

A-Archenteron
AMs-Aboral muscles
CB-Cell body
CBN-Ciliary band neuron
CP-Coelomic pouch
FM-Fertilization membrane
GV-Germinal vesicle
HG- Hind gut
Hpf-Hours post fertilization
Inv-Invaginations
LG-Lateral ganglia
MC-Mucous cell
MeCs-Mesenchymal cells
Oe-Oesophagus
OeMs –Oesophageal muscles
OG-Oral ganglia
OH-Oral hood
PB-Polar bodies
PCs-Pyramidal cells
Post-Post-oral ciliary band
Pre-Pre-oral ciliary band
Sc-Sensory cells
St-Stomach
Sts-Stomach sphincter
VP- Vegetal plate
